# Comparative analysis of CRISPR off-target activity discovery tools following *ex vivo* editing of CD34^+^ hematopoietic stem and progenitor cells

**DOI:** 10.1101/2022.09.09.507306

**Authors:** M. Kyle Cromer, Kiran R. Majeti, Garrett R. Rettig, Karthik Murugan, Gavin L. Kurgan, Jessica P. Hampton, Christopher A. Vakulskas, Mark A. Behlke, Matthew H. Porteus

## Abstract

While CRISPR-based editing most often occurs at DNA sequences with perfect homology to the guide RNA (gRNA), unintended editing can occur at highly homologous regions (i.e., off-target (OT) sites). Due to the pace at which genome editing therapies are approaching clinical applications, there is an emerging need to define effective workflows for investigating OT editing effects. A number of homology-dependent, *in silico-based* prediction methods and wet lab-based empirical methods exist to investigate OT editing, but few have been subjected to analytical assessment or head-to-head comparison in human primary cells using an *ex vivo* editing process optimized for high-fidelity gene editing. Therefore, we sought to compare publicly available *in silico* tools (COSMID, CCTop, and Cas-OFFinder) as well as empirical methods (CHANGE-Seq, CIRCLE-Seq, DISCOVER-Seq, GUIDE-Seq, and SITE-Seq) in the context of *ex vivo* hematopoietic stem and progenitor cell (HSPC) editing. To do so, we edited CD34^+^ HSPCs using 11 different guide RNAs (gRNAs) complexed with HiFi Cas9, then performed targeted next-generation sequencing of ~200-site panels containing a range of nominated OT sites identified by *in silico* and empirical methods. We identified an average of 0.45 OT sites per gRNA at an indel detection limit of 0.5%. This study confirmed the marked improvement in specificity with HiFi Cas9 compared to wild-type Cas9 without compromising on-target activity when delivered as an RNP. Additionally, all HiFi Cas9 OT sites using a standard 20nt gRNA were identified by all OT detection methods with one exception (SITE-seq did not identify an OT generated by an AAVS1 gRNA). This resulted in high sensitivity for the majority of OT nomination tools, however due to the large number of false positives called by most methods, *in silico*-based COSMID and empirical methods DISCOVER-Seq and GUIDE-Seq attained the highest positive predictive value. We did not find the empirical methods identified off-target sites that were not also identified by bioinformatic methods when delivered as an RNP complex. Finally, this study supports that refined bioinformatic algorithms could be developed that maintain both high sensitivity as well as positive predictive value which would enable more efficient identification of potential off-target sites without compromising a thorough examination for any given gRNA.

## Introduction

The CRISPR-Cas9 technology enables precision engineering of the genome with single base-pair resolution^1^. The site specificity of this system is imparted by a guide RNA (gRNA)—typically containing a 20 nucleotide (nt) spacer sequence—which couples with the Cas9 nuclease allowing it to scan the genome for a suitable protospacer adjacent motif (PAM), then bind and cleave sequences that display homology to the specific gRNA. While the location of the highest cleavage activity often occurs at the intended (on-target) site with perfect homology to the gRNA, activity at sites with lower degrees of homology may also occur^2–4^.

Following a DNA double-strand break (DSB) at both on- and off-target (OT) sites, the cell’s endogenous DNA repair machinery will either resolve the break using the non-homologous end joining (NHEJ) pathway which can result in inserted or deleted base pairs (indels) adjacent to the break site, or repair the break using homologous recombination where the sister chromosome or exogenous DNA donor is used as a repair template. If these DNA repair pathways are not successfully utilized, the cell will undergo cell cycle arrest due to the presence of an unresolved DSB^5^. In addition to these outcomes of DSB resolution, it is also possible that lower frequency events may occur such as translocations, inversions, large deletions, or chromothripsis^6^.

While several CRISPR-based therapies have entered the clinic^7,8^, there remains continued discussion of the most effective methodologies used to determine the location and frequency of the possible deleterious outcomes of unintended Cas9 activity. Currently, a range of tools and workflows have been developed to identify possible OT sites in the human genome. The first of these—*in silico-based* (bioinformatic) tools—use the specific gRNA of interest as input in order to return a list of potential OT sites with varying degrees of homology to the gRNA that may be screened for activity. While cleavage by the Cas9:gRNA ribonucleoprotein (RNP) complex is known to be largely homology-dependent, there is concern that purely homology-based prediction tools may miss some sites harboring bona fide OT activity. Additionally, the computational approaches to identify homology vary between *in silico* tools, leading to discrepancies in the sites identified. Furthermore, these computational OT detection tools primarily search a consensus reference genome and are thus unable to account genetic variation that could lead to differential Cas9 activity across patients.

As the CRISPR field matured, wet lab-based empirical approaches were developed to identify DSBs regardless of gRNA homology. Examples of these methods include CIRCLE-Seq^9^, GUIDE-Seq^10^, and SITE-Seq^11^, all of which tag or enrich for DSBs following delivery of Cas9 and gRNA (Supplemental Table 1). These studies reported a vast number of sites with a range of OT activity, some of which were missed by homology-based in silico prediction tools. While alarming, each of these empirical methods identified OT sites following delivery of Cas9 and gRNA to cell-free genomic DNA or immortalized cancer cell lines, which are known to harbor polyploidy, aneuploidy, tumorigenic SNPs, and dysfunctional DNA damage repair mechanisms^12–14^. In addition, the typically rapid doubling time of immortalized lines may impact OT activity profiles by providing excess unwound genomic DNA substrate on which Cas9 may bind and cleave. Similar to bioinformatic methods, the empirical methods are usually performed on a single cell line or genomic DNA from a single source and thus also do not fully examine the diverse genomes between patients.

In clinical *ex vivo* editing, Cas9:gRNA complex is delivered transiently to live, primary cells with functional DNA damage repair processes^15^. To improve the specificity of this process, high-fidelity variants of Cas9 (e.g., HiFi Cas9) have been developed^16,17^ and are being incorporated into the latest translational efforts^18–20^. Because this clinical workflow departs significantly from the context in which the original empirical methods were conducted, there is a need to compare the performance of both *in silico-* and empirical detection tools to determine whether true OT sites are currently being overlooked in the clinic. The ideal OT detection method should not only display high sensitivity (i.e., capturing the majority of sites displaying unintended OT editing activity), but have high positive predictive value as well (i.e., reporting as few false positive OT sites as possible).

To determine the performance of the existing methods, we designed a study to compare *in silico*-based and empirical methods in terms of their ability to predict Cas9 OT activity using transient delivery of high-fidelity Cas9:gRNA RNP to primary human hematopoietic stem and progenitor cells (HSPCs) *ex vivo*. First, we selected a series of 11 gRNAs previously investigated in the literature with a range of predicted activity at OT sites. From there, we compared the similarity and differences between OT sites nominated by various methods, and interrogated sites that were both unique as well as those that were largely overlapping across methods for evidence of OT editing in the *ex vivo* HSPC system. We then classified sites as true or false positives and evaluated the capability of these various methods to successfully recover bona fide OT sites. In doing so, this work provides clarity on how to successfully produce genome editing safety data with a focus on high-fidelity *ex vivo* genome editing systems.

## Results

### *Development of NGS panels to compare performance of* in silico *and empirical OT prediction methods*

To compare the performance of both *in silico* and empirical OT prediction methods, we chose to edit primary human CD34^+^-purified HSPCs using 11 different gRNAs previously reported in the literature (Table 1). We chose these gRNAs due to their disease relevance and/or inclusion in prior studies, including many from the original empirical OT detection publications (Supplemental Table 1). In order to maximize the number of detectable OT sites, we chose guides that display a range of expected OT activity (Supplemental Fig. 1a). The distribution of selected guides had predicted OT scores (higher = less predicted OT) ranging from 79 to 0 according to IDT’s CRISPR-Cas9 Guide RNA Design Checker tool, with a median predicted OT score of 21 (Supplemental Fig. 1b).

**Table 1:**
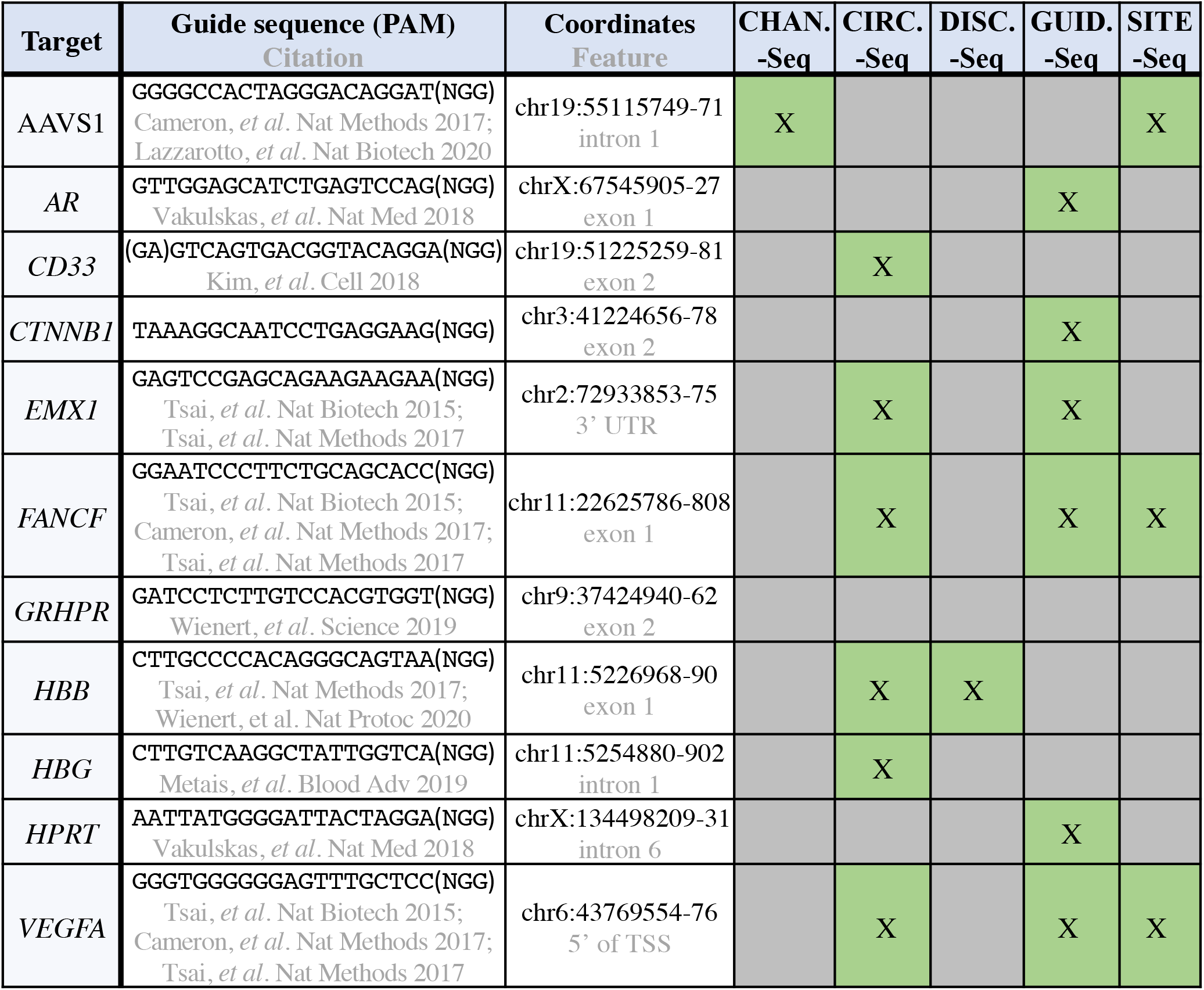
Summary of Cas9 gRNAs. Table depicts the characteristics of the gRNAs included in this study, including the gene name, coordinates (genome build hg38), and gene feature targeted by the gRNA. Green boxes denote inclusion in prior studies corresponding to the various empirical OT detection methods. Grey boxes indicate that data was unavailable for a given gRNA.

After choosing the gRNAs for this study, we then used three publicly available *in silico* prediction tools—COSMID^21^, CCTop^22^, and Cas-OFFinder^23^—to identify potential OT sites for each of the 11 guides. At the same time, we compiled all previously published OT data for the 11 gRNAs generated by empirical methods. These data were then used to establish custom 200-site panels for each gRNA that we interrogated using a rhAmpSeq-based next-generation sequencing (NGS) workflow—a standard method for identifying editing at potential OT sites^19^ (**Fig. 1a**).

**Figure 1:**
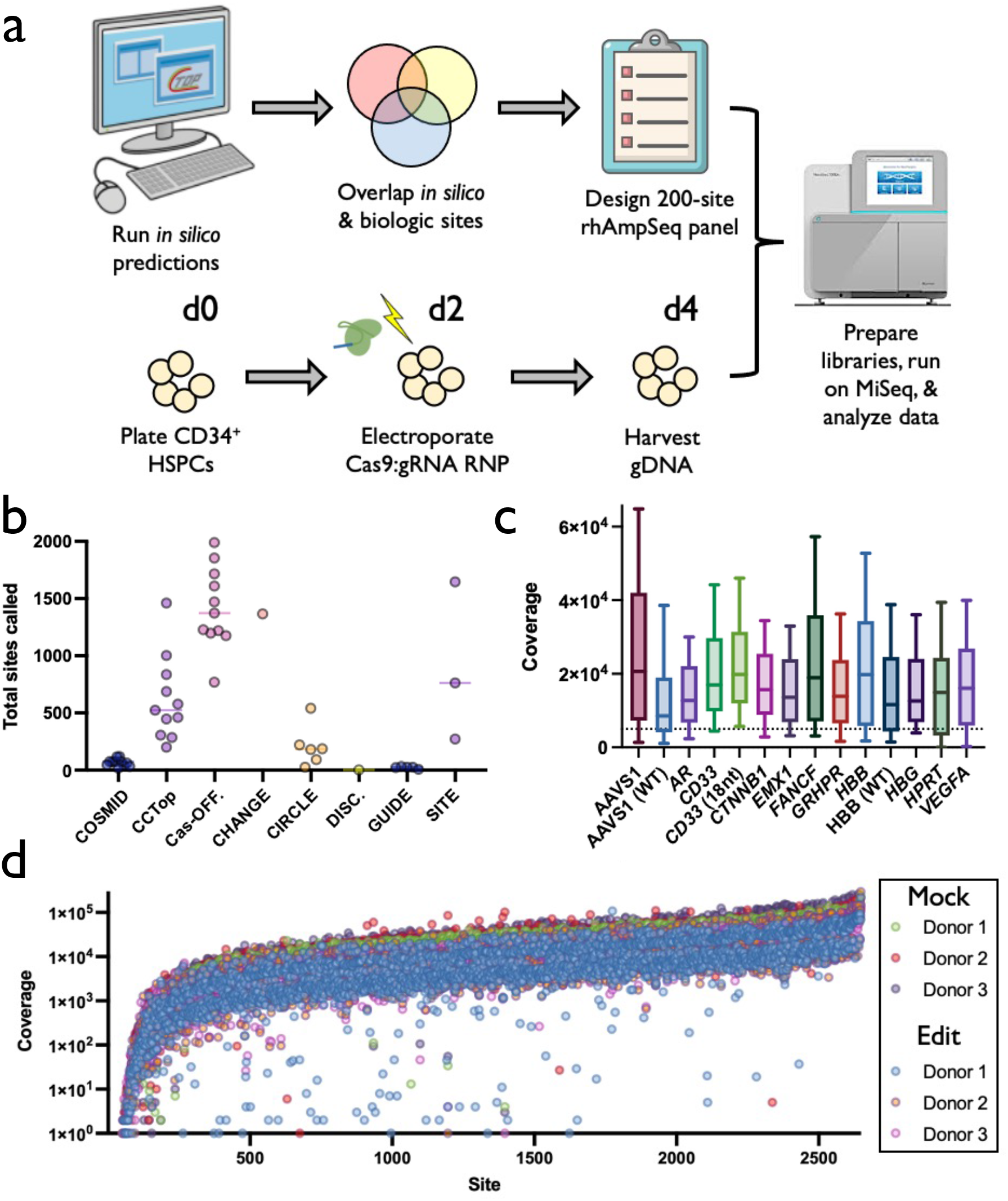
Experimental design & sequencing summary. a) Experimental design: *In silico* prediction programs (COSMID, CCTop, and Cas-OFFinder) were used to call potential OT sites for each gRNA. These sites were then overlapped with published data using empirical methods to discover high-likelihood sites of OT activity for each gRNA. Panels of 200 candidate sites were compiled and synthesized, which included those with high concordance across prediction methods as well as top-ranked sites called by individual methods. Concurrently, CD34^+^ HSPCs from 3 donors were edited via electroporation of Cas9 protein complexed with each respective gRNA and gDNA was harvested 2 days post-editing from mock and edited treatments. Libraries were prepared from gDNA, applied to the Illumina MiSeq platform, and NGS data was analyzed using a bioinformatic pipeline. b) Each dot depicts number of OT sites found for each gRNA by each discovery method. c) Coverage across all sites on panel for each gRNA (edited and mock treatments averaged at each site for all 3 HSPC donors). Middle line represents median, and box extends from 25^th^ to 75^th^ percentiles. Whiskers extend from 10^th^ to 90^th^ percentiles. Sequencing was performed in two separate rounds. All treatments were performed using HiFi Cas9 with 20-nt gRNA unless otherwise noted in parentheses. Dotted line represents 5,000x coverage. d) Each dots depicts read coverage at each individual OT site for a single donor. Shown on base 10 logarithmic scale.

We found that both *in silico* and empirical methods nominated a widely variable number of OT sites (**Fig. 1b**). While some of the tools returned a feasible number of nominated sites for follow-up screening (e.g., DISCOVER-Seq^24^, GUIDE-Seq, and COSMID found an average and standard deviation of 2.0+0.0, 22.6+9.1, and 66.4+32.2 sites per gRNA, respectively), others returned much larger lists of possible OT sites that would be time- and cost-intensive to screen by current methods (e.g., SITE-Seq, CHANGE-Seq^25^, and Cas-OFFinder found an average and standard deviation of 893.7+695.9, 1365.0+0.0, and 1418.4+353.6 sites per gRNA, respectively). In designing these 200-site panels for each gRNA, total number of sites nominated by each method was directly correlated with representation on each panel (i.e., more sites called by a given method, resulted in greater representation on each panel) (Supplemental Fig. 1c).

In the 200-site panels, we sought to not only include nominated sites that had a high degree of overlap across methods, but also those that were specifically detected by a single method and not by others (Supplemental Fig. 1d). We hypothesized that these two categories would allow us to identify the most likely loci with OT activity and determine whether a single method was capturing sites of bona fide activity that may have been missed by other prediction tools. Unsurprisingly, the tools that identified the largest number of candidate OT sites also had the highest degree of overlap with other detection methods (Supplemental Fig. 1e; note highest values in rows for Cas-OFFinder and SITE-Seq). The on-target editing site was also included in the rhAmpSeq panel.

### NGS panels reveal high on- and low-off-target editing profiles across all gRNAs

After designing the panels, we edited primary human CD34^+^ HSPCs by transiently delivering Cas9 RNP by electroporation—a workflow that has already been used to successfully treat patients suffering from β-thalassemia and sickle cell disease in the clinic^8^. We then harvested genomic DNA, prepared sequencing libraries, and sequenced each gRNA-specific panel on an Illumina MiSeq. We achieved an average coverage of 19,516 reads over all sites across all treatments, with 80.6% of sites exceeding our ideal coverage threshold of 5,000 reads to enable detection of low frequency editing events (**Fig. 1c**). We also observed a high degree of consistency for read counts at each site across donors and treatments (**Fig. 1d**). This indicated that the majority of sites were sequenced at high depth, while the regions prone to low coverage had consistently low read counts across all donors and treatments. We observed no apparent drop in read coverage in edited vs. mock treatments.

Following sequencing, we processed NGS data using CRISPAltRations (https://www.idtdna.com/pages/tools/rhampseq-crispr-analysis-tool)^26^. Analyzing the on-target site for each gRNA showed a high frequency of indels (median of 74.0% across all donors at all target loci) (**Fig. 2a & b**). Indel frequencies were also highly consistent across donors at each on-target site. While we identified high frequencies of editing at nearly every on-target site, we found that 94.8% of candidate OT sites (2,499 out of 2,635 total sites) had <0.1% indels after subtracting the mock background from our edited treatments at each site (**Fig. 2c** & Supplemental Fig. 2).

**Figure 2:**
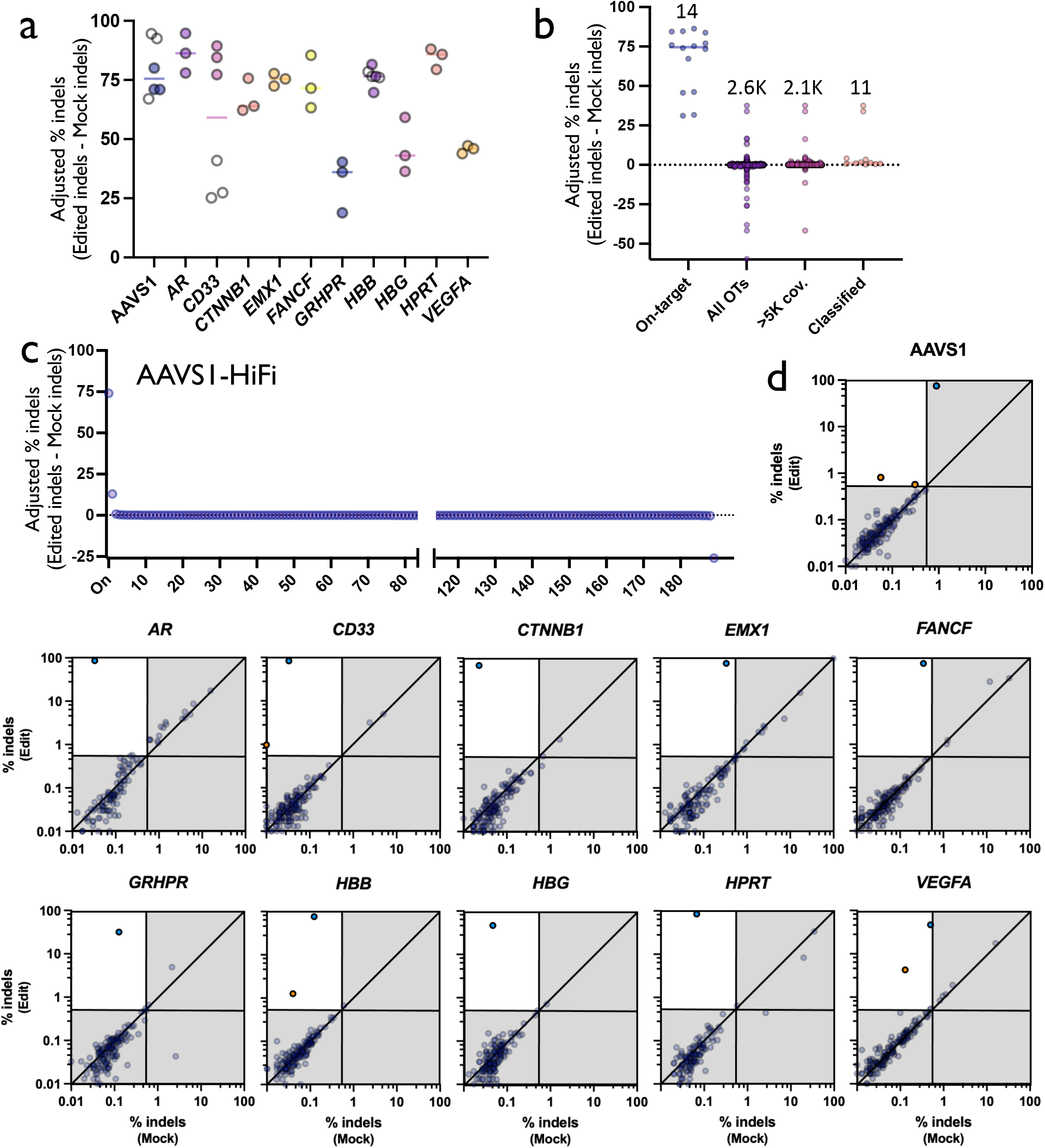
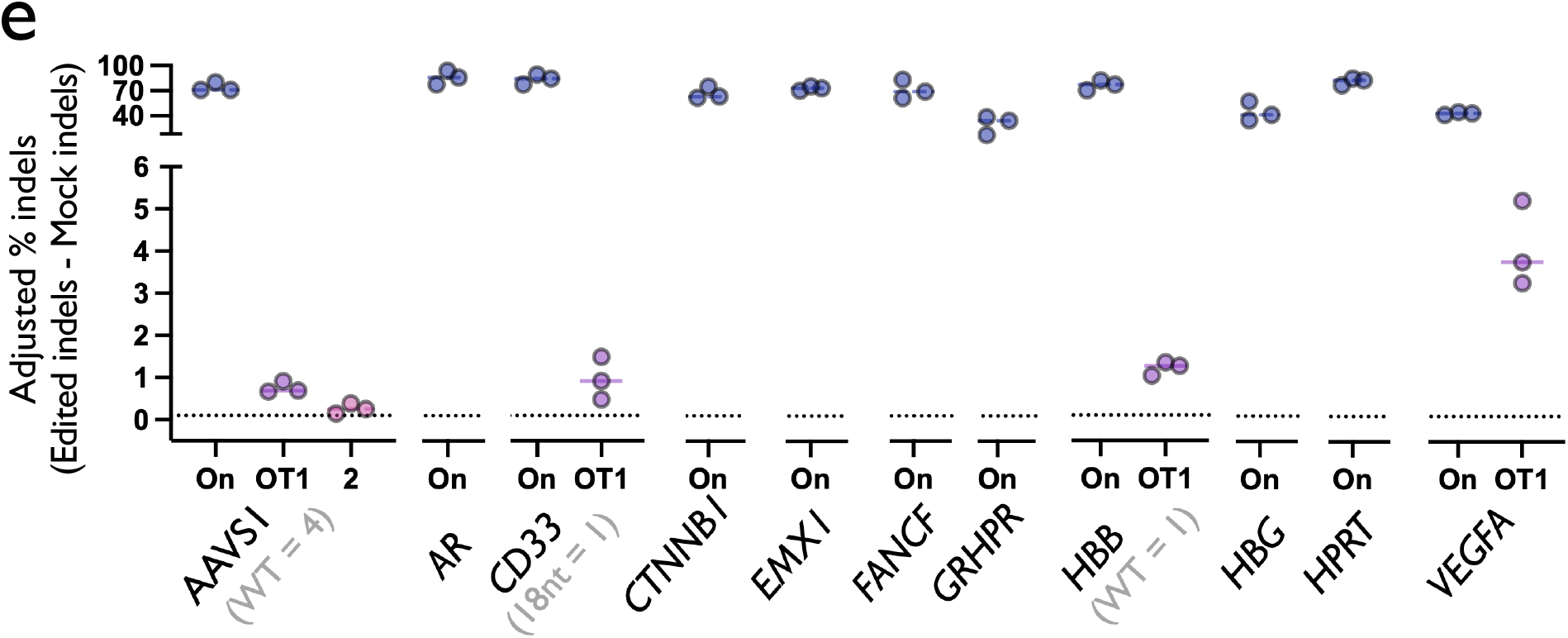
Summary of on- & off-target editing. a) On-target activity of each sgRNA determined by inclusion in rhAmpSeq NGS panel. White dots represent non-standard conditions (WT Cas9 treatment groups for AAVS1 and *HBB* gRNAs and truncated spacer for *CD33* gRNA), all others are edited using HiFi Cas9 with 20-nt spacer gRNAs. N=3 separate HSPC donors per treatment. b) Each dot depicts % adjusted indels (Edit-Mock) for each donor at the on-target as well as each OT site in each category. Numbers above dots indicate the total number of sites in each category. c) Each dot depicts avg. % indels (Edit-Mock) across donors at each site on panel in AAVS1-HiFi treatment (including the on-target site). Dotted line depicts 0.1% adjusted indel detection threshold after Mock is subtracted from Edited treatments. All sites are shown prior to filtering. d) Each dot depicts % indels averaged across donors for each site on panel with average coverage >5,000x. Top left quadrant indicates indels >0.5% indels in Edit treatment and <0.4% indels in Mock treatment. Blue dots represent on-target indels and orange dots represent classified OT sites. Shown on base 10 logarithmic scale. e) Each dot depicts % indels (Edit-Mock) for each donor at the on-target as well as each OT site that remained post-filtering using HiFi Cas9 and 20-nt spacers. Solid bars depict median % indels for all three HSPC donors. Grey text in parentheses indicates how many OT sites identified in alternative treatments. Dotted line depicts 0.1% adjusted indel detection threshold after Mock is subtracted from Edited treatments.

We then applied a binary classification method used previously (i.e., that determines presence or absence of editing) to determine if a nominated OT site is edited within a limit of detection of 0.5% indels given certain experimental bounds^26,27^. We found that the majority of sites probed met our coverage criteria of >5,000x (2,118 out of 2,635 total sites) (**Fig. 2b**). However, 99.5% of sites with adequate coverage displayed indels frequencies below our limit of detection (2,107 out of 2,118 total sites)— yielding only 11 total OT sites for classification (**Fig. 2d & e**). Importantly, all 11 sites were found to have p-values <0.05 by Fisher’s exact test. The majority of indels skewed toward deletions rather than insertions (Supplemental Fig. 4), and indel spectrums were unique to each gRNA in a fashion highly consistent across HSPC donors. Remarkably, the total number of on-target editing events exceeded the total number of classified OT editing events across all conditions (14 vs. 11 sites, respectively).

### Comparison to non-standard editing conditions reveals HiFi Cas9 dramatically reduces OT editing

When editing with HiFi Cas9 and a 20-nt gRNA, we found that the majority of gRNAs tested in this study showed no evidence of OT activity. In fact, 4 out of the 11 bona fide OT sites identified using non-standard conditions (3 from WT Cas9 treatments and 1 from truncated gRNA treatment) had no measurable indels when using HiFi Cas9 with a 20-nt gRNA. Compared to WT Cas9, the use of HiFi reduced the total number of detectable OT events in the AAVS1 treatment from 4 to 2 total OTs. We also found that HiFi Cas9 dramatically reduced the frequency of OT editing at the top OT site by an average of 36.8-fold compared to WT Cas9 using AAVS1 and *HBB* gRNAs, without compromising on-target editing frequency (**Fig. 3a & b**). And while we found that a previously reported 18-nt spacer gRNA targeting *CD33*^28^ appeared to reduce OT activity, it is difficult to determine whether this is a function of the truncated gRNA being more specific or simply possessing less total activity altogether (indicated by the lower on-target editing frequency of the shorter gRNA, an average of 31.2% vs. 83.8%) (**Fig. 3c & d**). In these comparative treatments, we observed a high degree of consistency in the indel spectrums generated that appeared to be more dependent on the core gRNA sequence rather than the use of WT or HiFi Cas9 or an 18- vs. 20-nt spacer (Supplemental Fig. 4).

**Figure 3:**
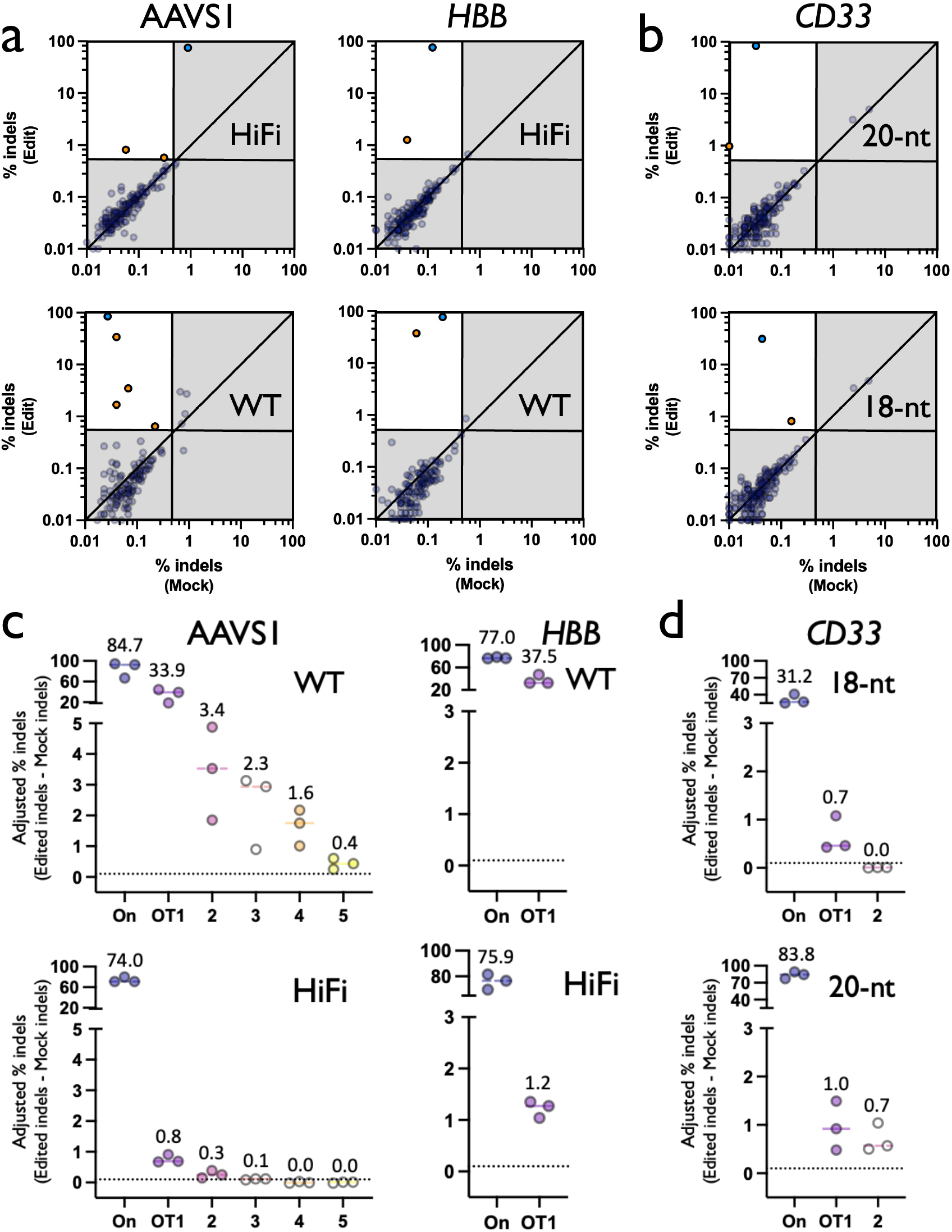
OT activity across comparative treatments. a) Each dot depicts % indels averaged across donors for each site on panel with average coverage >5,000x. Top and bottom panel represent treatments with HiFi and WT Cas9, respectively. Top left quadrant indicates indels >0.5% indels in Edit treatment and <0.4% indels in Mock treatment. Blue dots represent on-target indels and orange dots represent classified OT sites. Shown on base 10 logarithmic scale. b) Same as above, but with top and bottom panel representing treatments with 18-nt truncated and 20-nt *CD33* gRNAs, respectively. c) Each dot depicts % adjusted indels (Edit-Mock) for each donor at the on-target as well as each OT site that remained post-filtering. Top and bottom panel represent treatments with WT and HiFi Cas9, respectively. Solid bars depict median % indels for all three HSPC donors. Dotted line depicts 0.1% adjusted indel detection threshold after Mock is subtracted from Edited treatments. White dots represent OT sites that did not reproduce across treatments. d) Same as above, but with top and bottom panel representing treatments with 18-nt truncated and 20-nt *CD33* gRNAs, respectively.

### HiFi Cas9 OT sites for all selected gRNAs are called by most methods

Of the 11 true OT sites identified, we found that all were called by at least one *in silico* prediction method, but no method successfully called all 11 sites (**Fig. 4a**). However, under conditions of HiFi Cas9 and 20-nt gRNA, all 5 true OT sites were found by all *in silico* prediction methods. We also found that empirical methods reliably captured true OT activity as well, with only a single OT editing event missed by SITE-Seq in the AAVS1 treatment (which also happened to be the site with the lowest adjusted indel frequency—0.26%). For the 7 gRNA conditions with no detectable OT activity, all methods therefore reported no false negatives (**Fig. 4b**). Further investigation of our classified OT sites indicated that tolerance of gRNA mismatches increases with corresponding distance from the PAM site (**Fig. 4c**). We also did not observe any true OT sites that disrupted the NGG PAM used by SpCas9. Heavy dependence on the core guide sequence as well as an intact PAM has been well-documented previously^29^.

**Figure 4:**
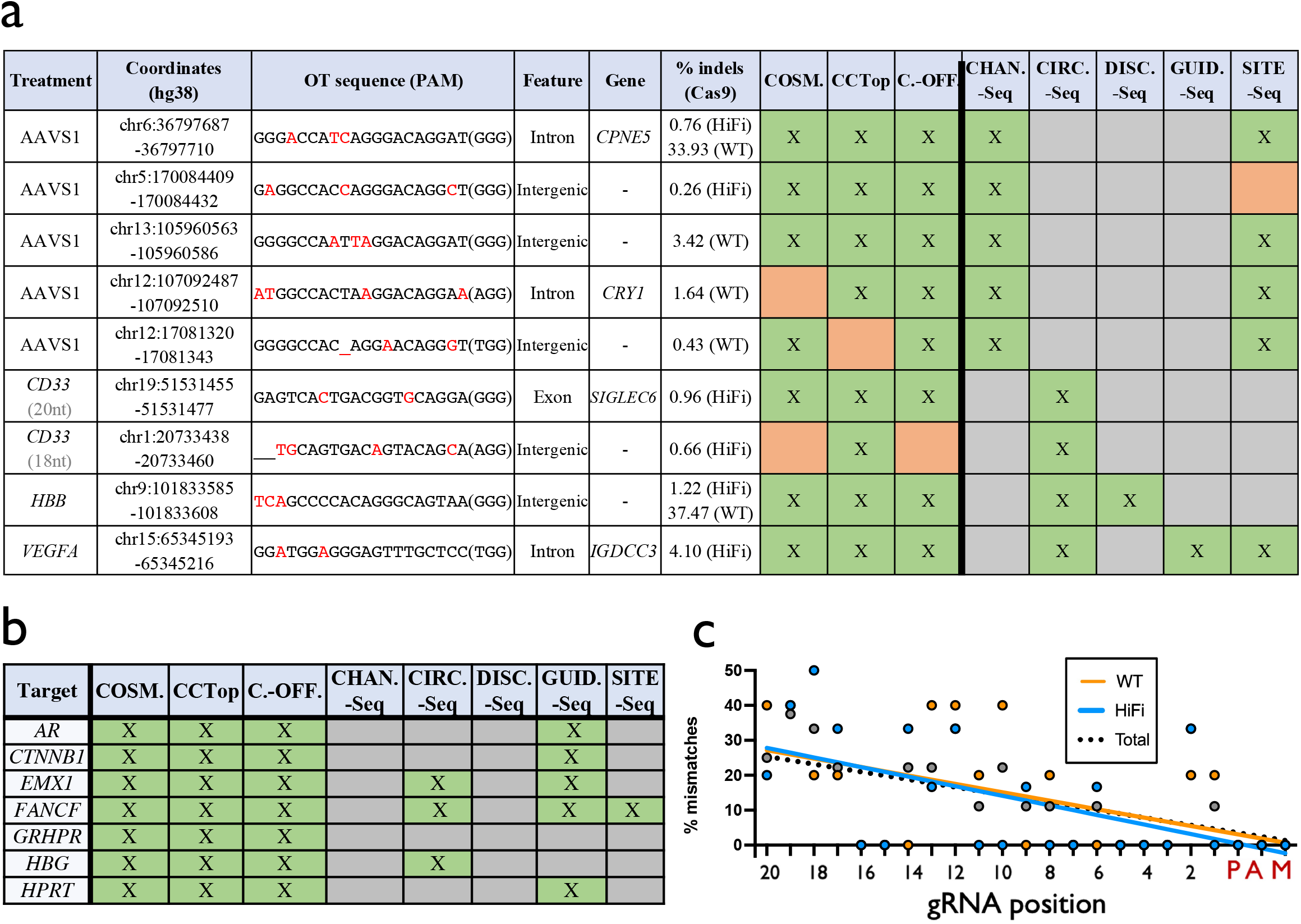
Summary of OT activity. a) Table depicts the characteristics of all true positive and false negative OT sites identified in this study, including gene name, coordinates, and gene feature targeted by the gRNA. % indels was calculated as % indels in Cas9-treated conditions, minus background % indels in Mock controls (averaged across all 3 donors for a given OT site). Green boxes denote successful prediction for a particular OT site by a particular detection tool. Orange boxes indicate a false negative for a particular OT site for a given detection tool. Grey boxes indicate that data was unavailable for a given gRNA and method. b) Table depicts all gRNAs that had no confirmed OT sites, and methods that therefore yielded no false negatives (indicated by green boxes). Grey boxes indicate that data was unavailable for a given gRNA and method. c) % mismatches at each position of the gRNA for 6 total OT sites in 11 HiFi Cas9 conditions and 5 total OT sites in 2 WT Cas9 conditions. Individual values and linear trendlines are plotted for each condition. Sites edited in both WT and HiFi treatments were only counted once in “Total” grouping.

We next sought to compare the performance of different OT detection tools by quantifying both sensitivity and positive predictive value (PPV) using our panel of sites that had sufficient coverage to perform binary editing classification (>5,000x coverage). For the sake of calculating PPV, any variants that were not highly ranked enough by a tool’s criteria to make our 200-target panels were considered a false positive. Due to the low number of true OT sites identified and the fact that most of these were captured by the majority of detection tools, we observed high sensitivity across all methods (**Fig. 5a**). When editing under standard conditions (HiFi Cas9 and 20-nt spacer gRNA), average sensitivity across all methods was 0.98, and all *in silico* methods attained sensitivity of 1.0. However, when sites were edited under more promiscuous, non-standard conditions (WT Cas9 or truncated gRNA), average sensitivity fell to 0.82. All empirical detection methods except SITE-Seq had a sensitivity of 1.0 for all guides that the method was performed on. SITE-Seq had a decreased sensitivity of 0.5 at the AAVS1 locus. However, there was a greater difference among methods when quantifying PPV (**Fig. 5b**). With the exception of DISCOVER-Seq that had a PPV of 0.5 (although data was only available for the *HBB* gRNA, and only nominated 2 sites), all other methods had PPVs <0.05 for all gRNAs tested. In other words, each method called far more false positives than true positives—though that could be expected due to the exceedingly low rate of OT activity generated in the *ex vivo* editing workflow.

**Figure 5:**
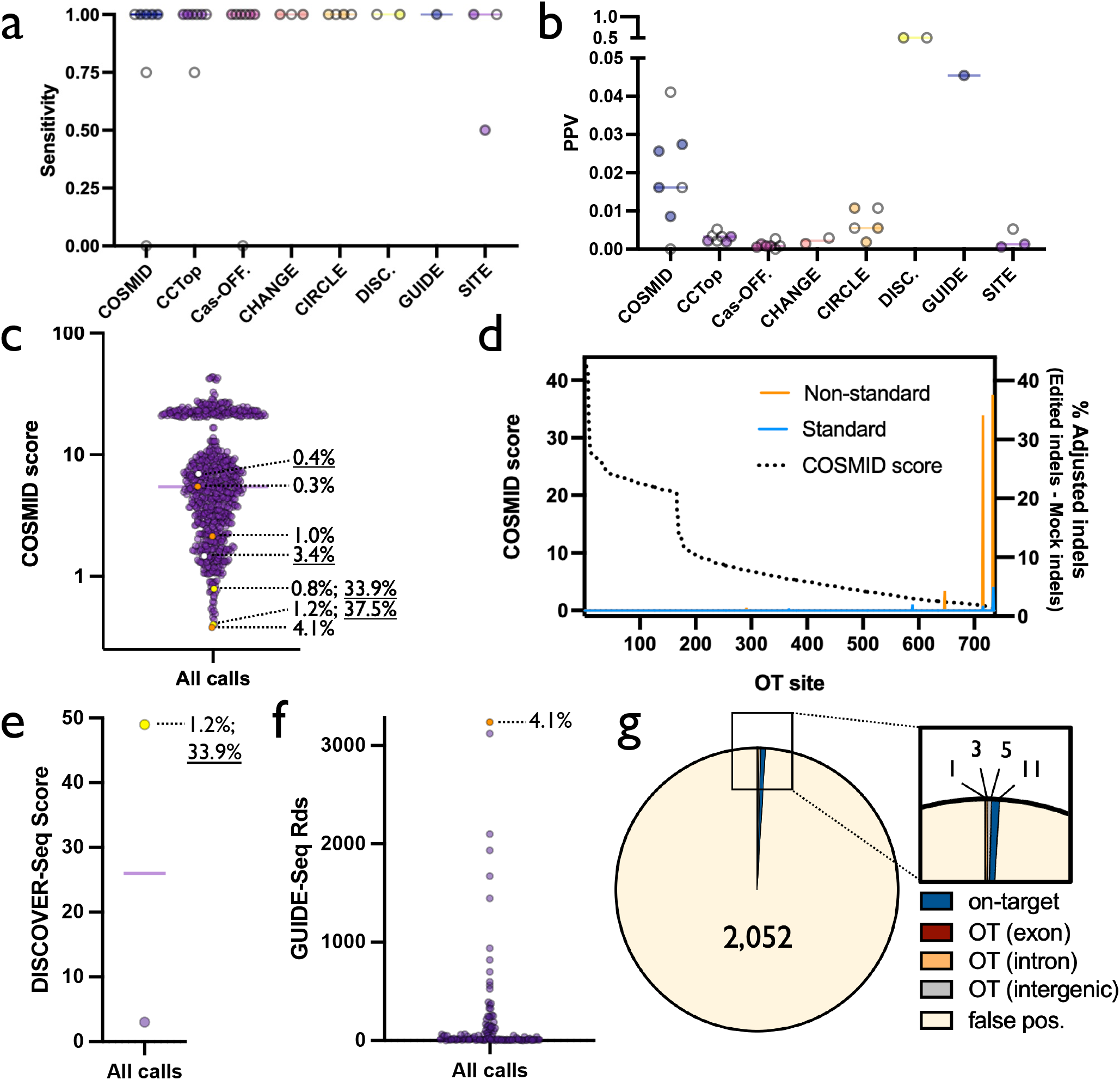
Sensitivity & specificity of each discovery method. a) Each dot depicts sensitivity for each gRNA for each discovery method. White dots indicate results derived from non-standard (i.e., WT Cas9 or truncated gRNA) conditions. Note: sensitivity was unable to be calculated in treatments where no OT sites were found. b) Each dot depicts positive predictive value (PPV) for each gRNA for each discovery method. All sites not on panel are assumed to be false positives. White dots indicate results derived from non-standard conditions. Note: PPV was not plotted for treatments where no OT sites were found. c) Each dots depicts COSMID score for all candidate OT sites for all gRNAs. True positives and corresponding indel frequencies are shown by dotted lines (WT indel frequency underlined). Orange, white, and yellow dots indicate OTs generated by HiFi, WT, and both HiFi and WT Cas9, respectively. d) For each OT site—rank-ordered left to right from high COSMID score to low along the x-axis—COSMID score and adjusted indel % (Edit-Mock) is plotted in standard and non-standard treatments. e) Each dot depicts the score assigned to a single DISCOVER-Seq OT site following editing using *HBB* gRNA. True positive and corresponding indel frequency is shown by dotted line (WT indel frequency underlined). Yellow dot indicates OT generated by WT and HiFi Cas9. f) Each dot depicts the number of reads covering a single GUIDE-Seq OT site following editing with *AR, CTNNB1, EMX1, FANCF, HPRT*, and *VEGFA* gRNAs. True positive and corresponding indel frequency is shown by dotted line. Orange dot indicates OT generated by HiFi Cas9. g) Pie chart summarizing proportion of on-target, OT, and false positives across all treatments for sites achieving sufficient coverage depth. False positives are defined as putative OT sites on panel that met coverage threshold that were not found to have OT activity.

When plotting all calls by COSMID (736 across all 11 intended gRNA cut sites), true OT sites were given likelihood-of-activity scores at a median in the 88^th^ percentile (in this case lower score indicates higher likelihood of OT activity) (**Fig. 5c**). While the vast majority of sites called did not reach our OT classification threshold for OT activity (727 out of 736 sites called), we found that a lower COSMID score both increased likelihood of OT activity as well as indel frequency should OT cleavage occur (**Fig. 5d**). In fact, the 2 top scores given to any OT site displayed OT activity, even with HiFi Cas9. We next plotted cumulative detection frequency as a function of COSMID score and found that 75% of all bona fide OT sites generated by standard conditions would have been found by interrogating the top 13% of all COSMID-nominated sites (Supplemental Fig. 4a). However, to achieve 100% detection, sequencing would have to have extended to the top 61% of all nominated sites.

While CCTop and Cas-OFFinder did not provide OT activity likelihood scores, we were able to group them into bins for number of mismatches and bulge sites from the target sequence (Supplemental Fig. 4b & c). For both methods, a greater number of sites were nominated as similarity to the gRNA decreased, and bona fide OT sites were clustered in bins with higher degrees of homology to the intended target. In assessing performance of the empirical methods, for DISCOVER-Seq and GUIDE-Seq—the two empirical methods with the highest degree of sensitivity and PPV—both true OT events occurred at the top-ranked call (determined by read count) by each method (**Fig. 5e & f**). Even among the empirical methods with lower PPV, true OT sites generally ranked at or near the top for CHANGE-Seq, CIRCLE-Seq, and SITE-Seq (Supplemental Fig. 4d-g).

### True OT sites predominantly map to non-coding regions of the genome

Overall, we found over two orders of magnitude more false positives than true OT sites within our NGS panels (**Fig. 5g**). Even though we also found more on-target activity than OT activity (11 on-target events vs. 9 OT events (including standard and non-standard conditions)), we nevertheless sought to determine whether these true OT sites may be likely to cause genotoxicity or an oncogenic expansion event. Of bona fide OT sites for our selected gRNAs, 5 out of 9 resided in intergenic regions of the genome, the effects of which remain difficult to interpret (**Fig. 4a & 5g**). Three of 9 bona fide OT sites resided in intronic regions of genes, indels in which do not typically disrupt exon splicing or gene expression^30^. Only a single OT site was found in the exon of a gene, *SIGLEC6*, which was caused by the *CD33* gRNA. The indel spectrum created by the *CD33* gRNA is consistent with a frameshift 1-bp insertion, which would likely knock out this gene (Supplemental Fig. 3). However, the detected indel frequency was <1% and knockdown of this gene is not known to impart oncogenic potential^31^. To better understand the overall likelihood that exons are OTs of gRNAs targeting annotated genes, we performed an *in silico* aggregated search for all possible on-target gRNAs targeting 19,222 human genes using the IDT CRISPR-Cas9 Guide RNA Design Checker tool and investigated the relative frequency that exons are nominated OTs. From this we found that an average of 7.2% ± 2.9% of predicted OTs for gRNAs targeting a gene are annotated as within exonic regions (Supplemental Fig. 5). This is comparable to our experimental findings here that 1 of 9 unique bona fide OTs target an exon. Exons only make up ~3% of total genomic content, so this is enriched approximately 2-fold compared to what could be expected if genomic regions were selected at random.

## Discussion

In this study, we investigated the efficacy of different OT nomination tools to successfully identify sites with a high likelihood of Cas9 activity in order to inform genome editing safety investigations. When editing HSPCs in a high-fidelity *ex vivo* genome editing workflow, we found that out of 2,061 potential OT sites identified by both *in silico* and empirical methods from 11 different previously identified targets, there were only 11 sites with detectable indels when the Cas9 nuclease is delivered as an RNP into repair-competent healthy human HSPCs. Importantly, these numbers only include sites on our NGS panels that reached sufficient coverage depth and does not count the greater number of lower-ranked sites called by each method that were not included on our panels. In addition, supporting previously published data^16^, the use of HiFi Cas9 resulted in a >30-fold reduction in the frequency of off-target indels without a reduction in on-target activity. These results demonstrate that the CRISPR-Cas9 system can be utilized with very high specificity. The results shown an average of <1 detectable OT event per gRNA to a limit of detection of 0.5% indels across a set of guides with highly variable predicted OT severity. We also found that existing OT detection tools, both *in silico* prediction and empirical methods, identify the majority of detectable OT events for the guides interrogated, especially using high-fidelity genome editing conditions (HiFi Cas9 delivered as an RNP). Notably all of the bioinformatic programs identified true OT sites when editing with HiFi Cas9.

We believe that our results contrast with prior findings for several reasons. First, we delivered Cas9 to primary cells rather than immortalized cell lines that often harbor gross chromosomal abnormalities (polyploidy, aneuploidy, translocations, etc.) with dysfunctional DNA damage and nucleic acid delivery-sensing responses^32–34^. Furthermore, delivery of Cas9 to living cells likely serves as a better model for Cas9 binding and cleavage due to the presence of vast regions of inaccessible chromatin^35^ as well as active DNA damage repair mechanisms. These biological phenomena are likely contributing factors for our finding that methods relying on delivery of Cas9 to cell-free genomic DNA—CHANGE-Seq, CIRCLE-Seq, and SITE-Seq—attained the lowest PPVs of all empirical methods tested (**Fig. 5b**). Additionally, unlike prior studies that relied on a pre-existing pool of cells with oncogenic mutations^36,37^, by conducting our experiments in primary HSPCs that do not harbor such aberrations^38^, it is not surprising that this study found no bona fide OT sites residing in exons of known tumor suppressors or oncogenes. This was further confirmed by a recent publication that found no evidence that Cas9 introduced or enriched for oncogenic mutations following *ex vivo* editing in primary HSPCs^38^. Taken together, our study highlights the importance of evaluating Cas9 OT detection tool performance in the appropriate application-specific context (e.g., high-fidelity nuclease, primary HSPCs) where epigenetic factors and DNA damage repair mechanisms can impact the appearance of unwanted OT editing.

Collectively, our findings reinforce the importance of using high-fidelity variants of Cas9 to dramatically reduce the frequency of unintended editing events. Since HiFi Cas9^16^ reduces editing at OT sites without compromising on-target editing, it is also likely that the risk of translocations/inversions between the on-target and OT sites are reduced accordingly and has been shown in one example^19^. Furthermore, other high-fidelity Cas9 enzymes as well as gRNA innovations have been reported^17,39,40^, and continued efforts attempt to increase activity and specificity of Cas9 via protein engineering. This may be especially important as genome editing technology begins to be adapted for *in vivo* delivery^7^.

While this study did not detect a large number of OT events, the experimental workflow primarily allowed us to capture the most common unintended genomic event following Cas9 editing—small, sitespecific indels at sites other than the targeted locus. Other abnormalities such as translocations, inversions, large deletions, or chromothripsis would have likely been overlooked by our methodology. In fact, chromothripsis has even been documented as a consequence of on-target Cas9 cleavage^6^, a phenomenon our experimental workflow cannot capture accurately. While these other abnormalities are reported to occur at relatively low frequencies, further studies are needed to better understand, and possibly prevent, creation and enrichment of cells harboring large-scale genomic disruptions following CRISPR-based editing.

Our study provides a comparison of OT detection tools in the context of genome editing HSPCs using high-fidelity nucleases. These tools approach the problem of finding Cas9-generated OT edits from two different directions—*a priori* or *a posteriori*. The *a priori in silico* prediction tools operate under the assumption that Cas9 binding and cleavage requires presence of the PAM and homology to the gRNA, while *a posteriori* empirical methods identify OT sites experimentally. However, OT sites nominated by empirical methods are often so numerous that filtering based on homology is often used as a quality control step^41^. Therefore, it is reasonable to ask, if filtering based on homology to the intended cut site is recommended during processing putative OT sites identified by empirical methods, then how do we interpret sites that pass coverage metrics but display no homology to the gRNA? Despite deliberate inclusion of OT sites on NGS panels that were nominated only by empirical tools, we found none of these sites to harbor detectable levels of OT editing, therefore leading to the conclusion that minimal Cas9 activity occurs at sites lacking homology to the gRNA.

While *in silico*-based prediction of OT sites requires minimal time, effort, and expertise, empirical methods are comparably cumbersome to establish in the lab and require the development of specialized skills to be used robustly and reproducibly. Though datasets for empirical methods were not available for many of the guides used in this study, we found no OT sites validated under high-fidelity editing conditions identified solely by empirical methods. While *in silico* methods appear capable of capturing most true OT events, we note that they rely on homology to a consensus genome and therefore would not take into account patient-specific variation, which has been shown to impact Cas9 activity^38^. The empirical methods, however, are also limited by not being performed on a diverse array of genomes and are usually also performed on a single genome. As a solution, both *in silico* and empirical methods could be performed on diverse genomes, especially *in silico* methods as the catalogue of sequenced human genomes continually grows.

Taken together, our results provide a comparative analysis of the relative performance of currently available OT detection methods. While no one method emerged as ideal for detecting all OT events without a high frequency of false positives, certain methods had a better profile of sensitivity and positive prediction value than others, particularly using a high-fidelity system. COSMID, for example, is an easily accessible online program that requires no user expertise and identified all of the bona fide OT sites using the high-fidelity system while having the highest positive prediction frequency of *in silico* programs investigated. All of the bioinformatic tools evaluated are several years old, however, and it is likely that improved tools using machine learning or other techniques could be developed in the future using datasets like these and others^29^. For example, while we could not define a clear cut-off for *in silico* tools like CCTop or Cas-OFFinder which do not incorporate a score, the bona fide OT sites were most likely to occur at sites with the greatest degree of homology. The next generation of *in silico* algorithms, therefore, should be able to incorporate a score (as COSMID does) into the tool to help investigators determine how to prioritize analysis of OT sites. The efficient characterization of genomic outcomes of Cas9 activity, both intended and not, will help ensure that genome editing-based therapies are developed and delivered in a way that maximizes safety and minimizes the risk of genotoxic or oligoclonal expansion events while facilitating the translational development of these potentially powerful therapies.

## Methods

### Acquisition of HSPCs

Primary human HSPCs were sourced from fresh umbilical cord blood (generously provided by Binns Family program for Cord Blood Research) under protocol 33818, which was approved and renewed annually by the NHLBI Institutional Review Board (IRB) committee. All patients provided informed consent for the study. Patient information was de-identified prior to laboratory experiments—we therefore are unable to make a statement speaking to sex or ethnicity of participants. Donors were not aware of the research purpose or compensated for their participation. Consent forms provided express permission to publish de-identified genetic information.

### *Ex vivo* culturing of HSPCs

CD34^+^ HSPCs were bead-enriched using Human CD34 Microbead Kits (Mitenyi Biotec, Inc., Bergisch Gladbach, Germany) according to manufacturer’s protocol and cultured at 1×10^5^ cells/mL in CellGenix GMP SCGM serum-free base media (Sartorius CellGenix GmbH, Freiburg, Germany) supplemented with stem cell factor (SCF)(100ng/mL), thrombopoietin (TPO)(100ng/mL), FLT3–ligand (100ng/mL), IL-6 (100ng/mL), UM171 (35nM), 20mg/mL streptomycin, and 20U/mL penicillin.

### Genome editing of HSPCs

Chemically modified gRNAs used to edit HSPCs were purchased from Synthego (Menlo Park, CA, USA). The gRNA modifications added were the 2′-O-methyl-3′-phosphorothioate at the three terminal nucleotides of the 5′ and 3′ ends^39^. All Cas9 protein (both Alt-R^®^ S.p. HiFi and WT Cas9 nuclease) was purchased from IDT, Inc. (Coralville, IA, USA). HSPCs were cultured in vitro for 2d and then edited as follows: RNPs were complexed at a Cas9:gRNA molar ratio of 1:2.5 at 25°C for 10min prior to electroporation, HSPCs were resuspended in P3 buffer (Lonza, Basel, Switzerland) with complexed RNPs, and electroporated using the Lonza 4D Nucleofector (program DZ-100). Cells were plated at 1×10^5^ cells/mL following electroporation in the cytokine-supplemented media described above.

### rhAmpSeq panel design

The primary goal of rhAmpSeq panel design was to assemble a list of 200 sites with either a high degree of overlap across methods (representing sites of high likelihood of OT activity) or those that were found only by one or more empirical methods (representing sites that can inform whether purely homology-based prediction is missing true OT sites). To assemble these panels, we took virtually all sites with a high degree of overlap. Then for lower-overlap samples, we established a ranking system that allowed us to prioritize OT sites that were nominated by one or several methods. For COSMID, we used the internal score that is generated as output (lower score indicates higher likelihood of activity). For CCTop and Cas-OFFinder, where scores were not generated, we prioritized sites with the fewest mismatches and bulges. For DISCOVER-Seq, we were able to include both predicted OT locations on our panel. And for all other empirical methods, we prioritized the sites with the highest read counts from the highest concentration Cas9 conditions.

### rhAmpSeq library preparation

On- and off-target editing rates in HSPCs were measured by amplicon-based NGS. Genomic DNA from edited and unedited HSPC samples was harvested 48h post-editing using QuickExtract DNA Extraction Solution according to manufacturer’s recommendations (Lucigen Corp., Teddington, UK) and diluted to 4.55 ng/uL in IDTE pH 8.0 (IDT, Inc. (Coralville, IA, USA). Amplicon libraries were generated using target-specific rhAmpSeq primer panels (as described above) with 4x rhAmpSeq library mix 1 and 50 ng of gDNA input. The following cycling conditions were used for PCR 1: 95 °C for 10 min; [95 °C for 15 s; 61 °C for 8 min] × 14 cycles; 99.5 °C for 15 min; 4 °C hold. Target-specific amplicon libraries from PCR 1 were diluted 20-fold in nuclease-free water and were subsequently tagged with P5 and P7 Illumina adapter primers with dual unique indices via a second round of PCR using 4x rhAmpSeq library mix 2. The following cycling conditions were used for PCR 2: 95°C for 3 min; [95°C for 15 s; 60°C for 30 s; 72°C for 30 s] × 24 cycles; 72°C for 1 min; 4°C hold. Libraries were purified using the AMPure XP system (Beckman Coulter, CA, USA), and quantified using Qubit 1X dsDNA HS Assay Kit (ThermoFisher Scientific, MA, USA) or qPCR before loading onto the Illumina MiSeq or NextSeq 500/550 platform (Illumina, CA, USA). Paired-end, 150-bp reads were sequenced using V2 or V2.5 chemistry.

### Data analysis

NGS data were analyzed and CRISPR editing quantified using CRISPAltRations with a default window parameter for Cas9 (8bp)^26^. The following filter was used to determine sites that had sufficient data and bounds to be binarily classified as “Edited” or “Not Edited”: 1) >5,000x read depth; 2) >0.5% indels in the “Edited” treatment; and 3) <0.4% indels in the “Mock” treatment. These cutoffs were based on average values across Mock and Edited samples for all 3 HSPC donors. We also ensured that each OT site remaining post-filtering had at least 1 donor that met all coverage and indel frequency criteria on their own. To quantify sensitivity and PPV, the following values were determined for all treatment conditions, using the following definitions: 1) true positives = all sites called by a given method that showed detectable OT activity; 2) false positives = all sites called by a given method that met coverage metrics but remained below detection; and 3) false negatives = all true OT sites missed by a given method (See Extended Data). Sensitivity was then defined as the number of true positives captured as a percentage of all true positives and false negatives. PPV was defined as the number of true positives captured as a percentage of all true positives and false positives. To predict the specificity of previously published gRNAs, all sequences were submitted to the IDT gRNA design checker (CRISPR-Cas9 guide RNA design checker | IDT (idtdna.com)) and OT scores recorded.

## Data Availability

High-throughput sequencing data generated for all rhAmpSeq panels will be uploaded to the NCBI Sequence Read Archive (SRA) submission. The filtered data for all figures in this study are provided in the Supplementary Information/Source Data file.

## Statistical analysis

Statistical analysis for binary classification of editing was performed on the treated/edited samples comparing to the untreated/unedited samples using a Fisher’s exact test (p<0.05) using set parameters for read depth (>5,000x) and % NHEJ (>0.5 % in treated samples, <0.4% in untreated samples) based on previously published methods^26^.

## Acknowledgements

This project was supported by the Food and Drug Administration (FDA) of the U.S. Department of Health and Human Services (HHS) as part of a financial assistance award [Center of Excellence in Regulatory Science and Innovation grant to University of California, San Francisco (UCSF) and Stanford University, U01FD005978] totaling $75,000 with 100 percent funded by FDA/HHS. The contents are those of the authors and do not necessarily represent the official views of, nor an endorsement, by FDA/HHS, or the U.S. Government. *Products and tools supplied by IDT are for research use only and not intended for diagnostic or therapeutic purposes. Purchaser and/or user are solely responsible for all decisions regarding the use of these products and any associated regulatory or legal obligations*.

## Author Contributions Statement

MKC and MHP supervised the project. MKC and MHP designed experiments. MKC, KRM, GRR, KM, JPH, GK, CAV, and MAB carried out experiments. MKC wrote the manuscript.

