## Supplemental Figures & Tables for "Comparative analysis of CRISPR off-target activity discovery tools following *ex vivo* editing of CD34^+^ hematopoietic stem and progenitor cells"

Supplemental Table 1: Summary of empirical Cas9 OT discovery methods

| Method | Journal | Year | Cas9 | Delivery method | Cell types |
|---|---|---|---|---|---|
| CHANGE-Seq | Nature Biotechnology | 2020 | WT | Cell-free RNP | human primary T cells, HEK293, & U2OS cell lines |
| CIRCLE-Seq | Nat Methods | 2017 | WT | Cell-free RNP | HEK293, U2OS, & PGP1 cell lines |
| DISCOVER-Seq | Science | 2019 | WT | RNP electroporated into cells | K562, human iPSC, & murine B16-F10 cell lines |
| GUIDE-Seq | Nature Biotechnology | 2015 | WT | RNP electroporated into cells | HEK293 cell line |
| SITE-Seq | Nature Methods | 2017 | WT | Cell-free RNP | HEK293, K562, & HeLa cell lines |

Confidential - Company Proprietary

Supplemental Figure 1: Overlap of *in silico* & empirical methods

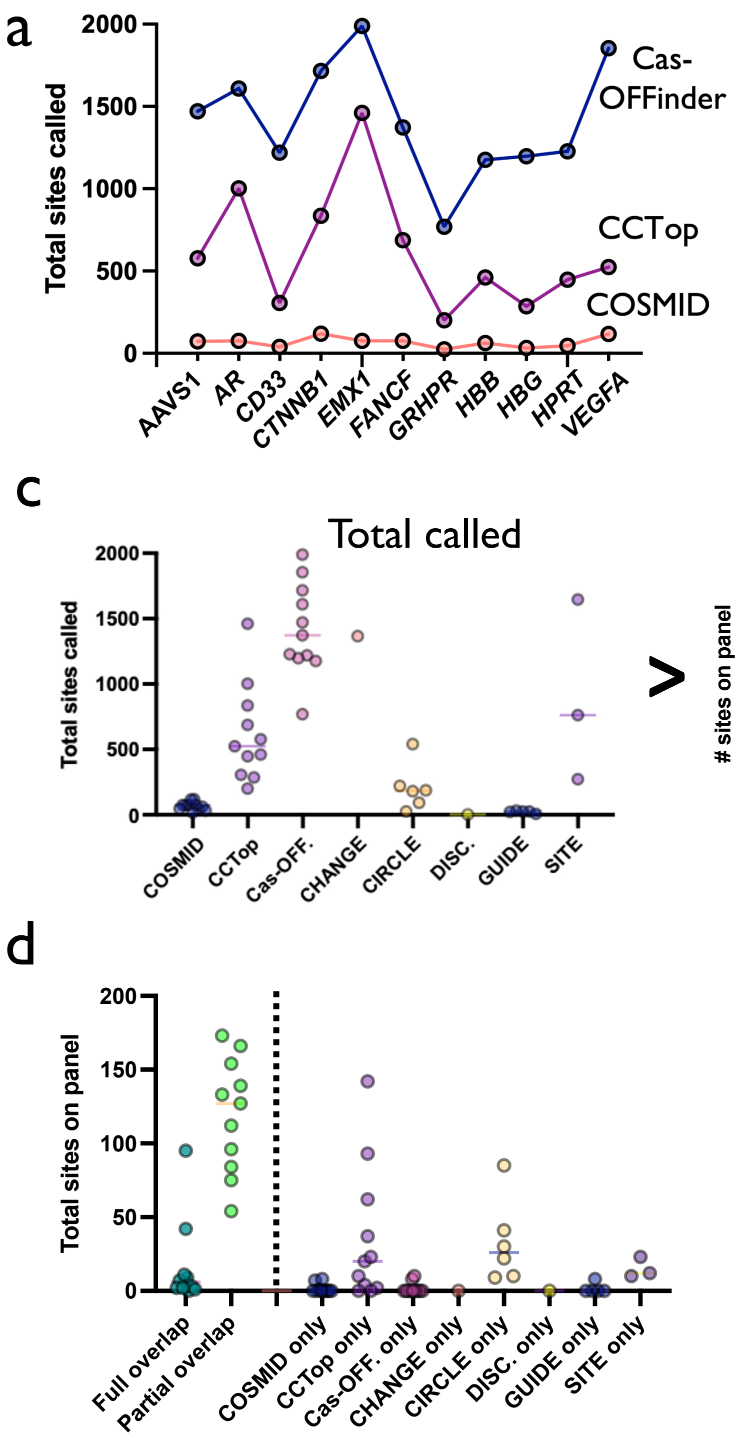

Supplemental Figure 1: Overlap of *in silico* & empirical methods

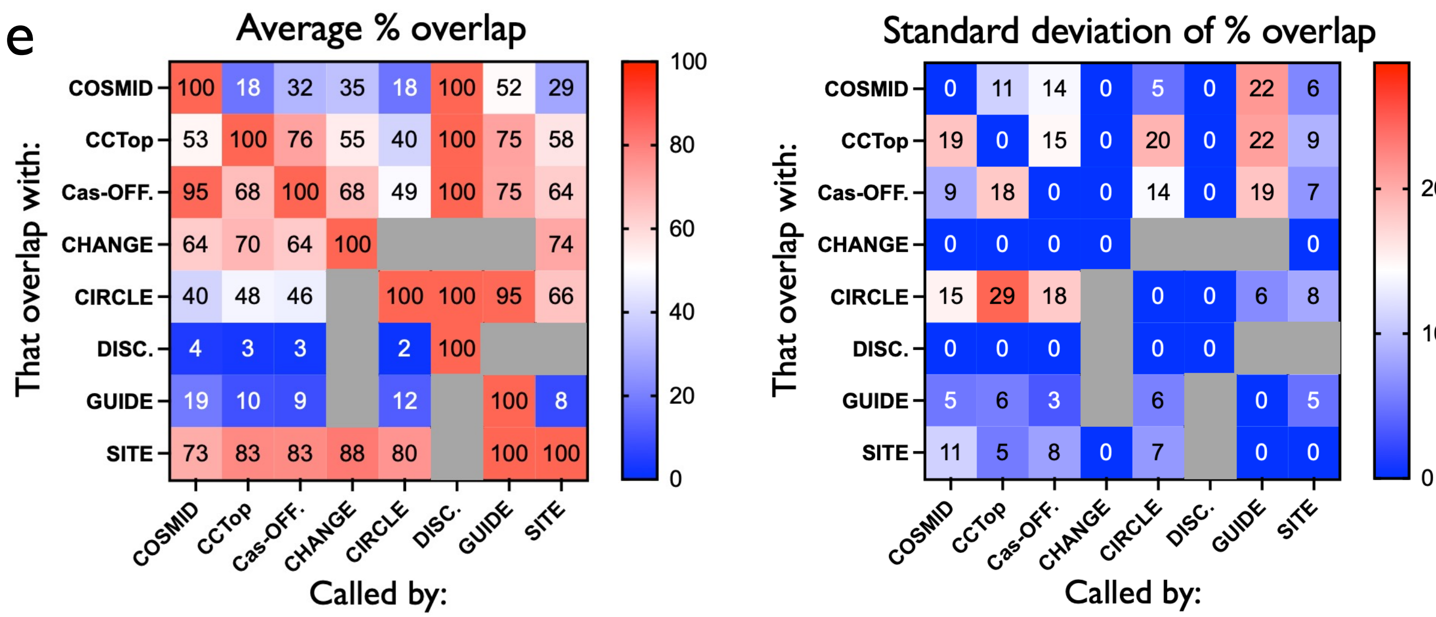


a) Each dot depicts number of OT sites found by each discovery method for each gRNA.
b) Predicted OT score by IDT gRNA assessment tool for each of 11 gRNAs used in this study.
c) Each dot depicts number of OT sites found for each gRNA by each discovery method. Left panel plots total sites called per gRNA for each method (identical to Fig. 1b, but displayed here for reference); right panel plots number of sites included on each gRNA panel for each method.
d) Each dot depicts number of sites on panel with full or partial overlap across all detection methods, as well as number of sites detected only by a single method (to the right of the dotted line).
e) Left panel shows heatmap depicting average number of OT sites per gRNA called by methods on the x-axis, which overlap with methods on the y-axis. Right panel depicts standard deviation of % overlap between methods.

Supplemental Figure 2: OT activity at each site on panel for each gRNA

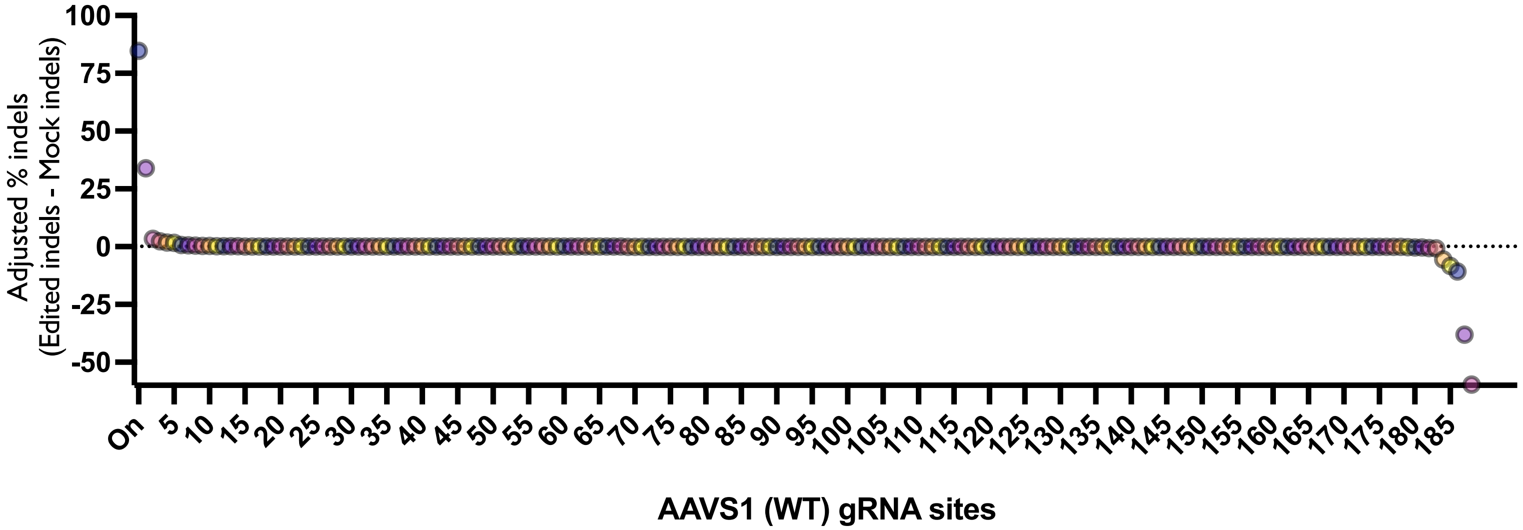


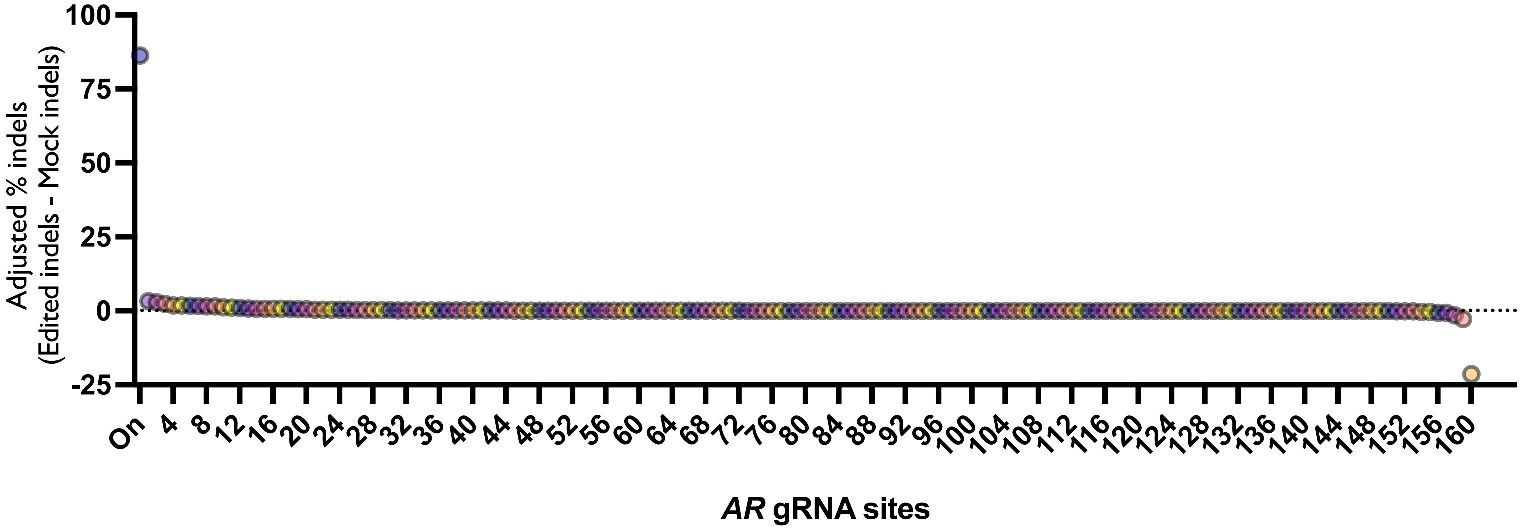

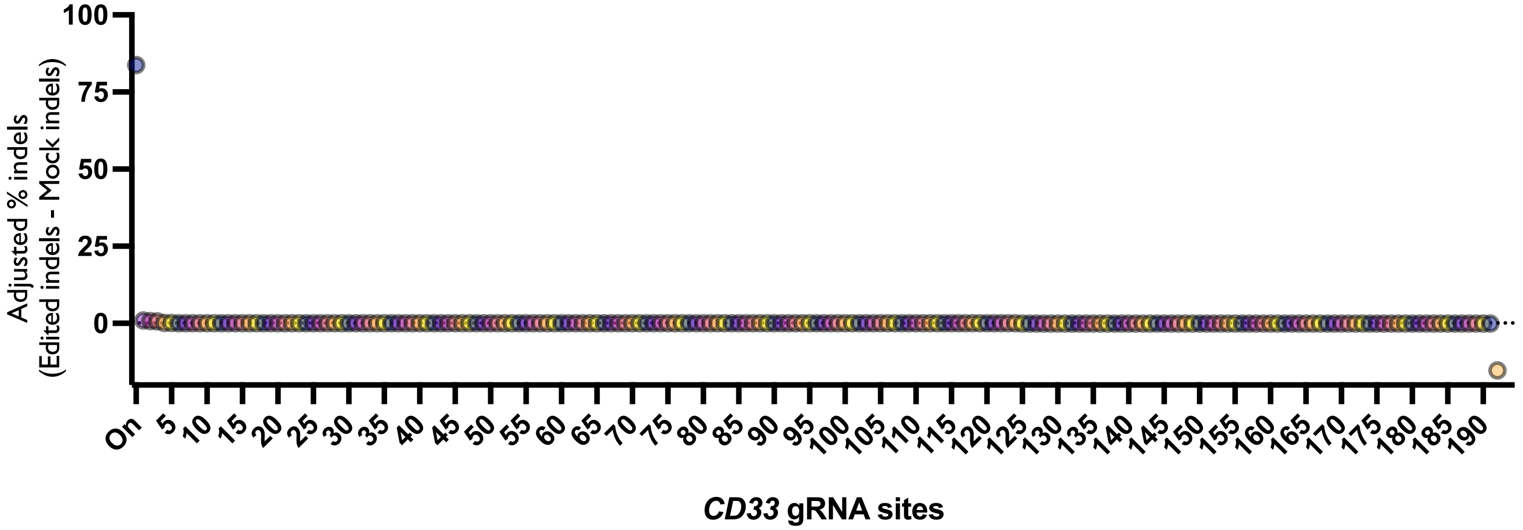


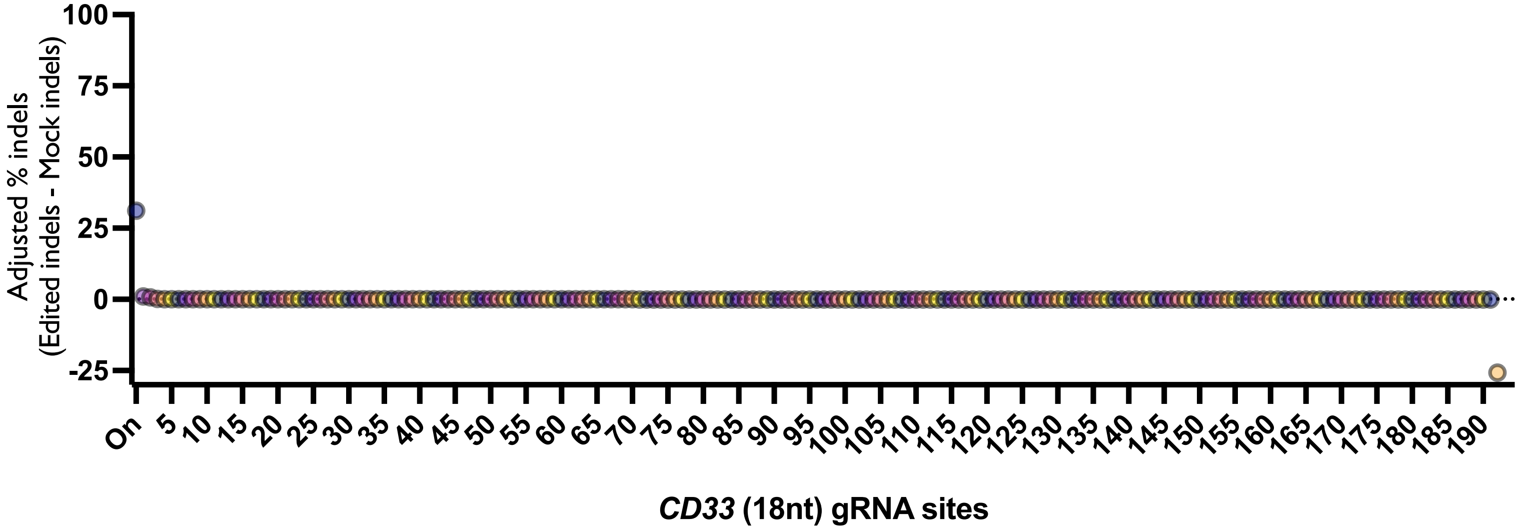


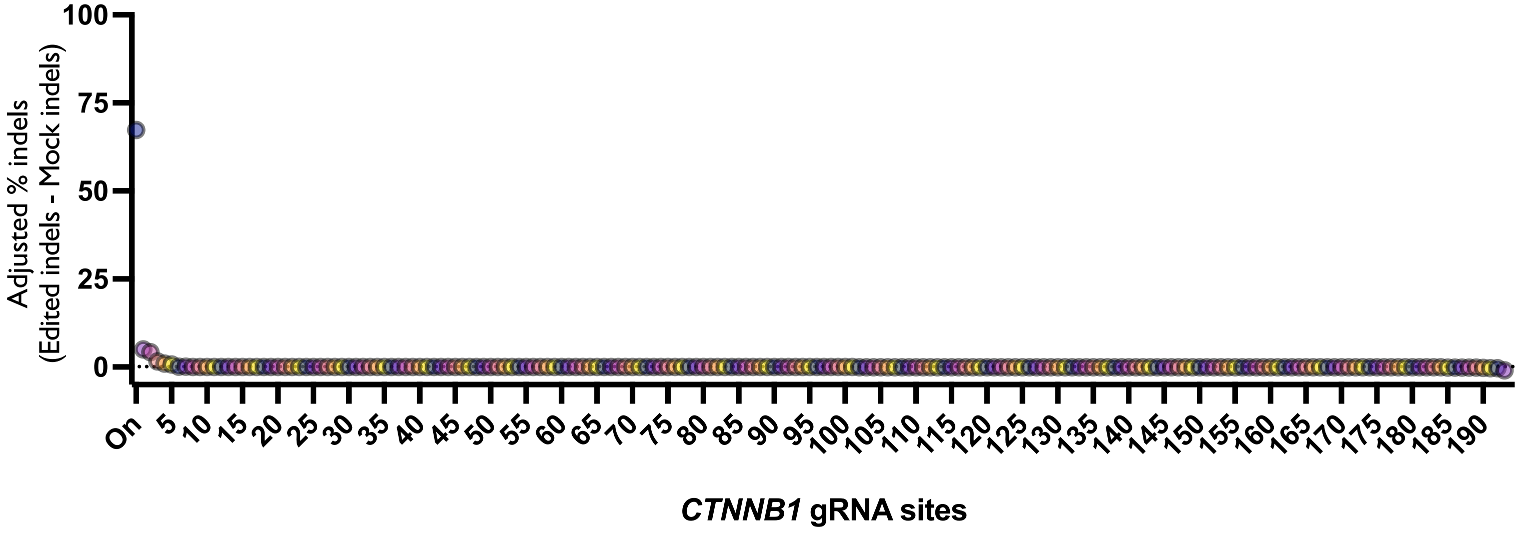


Confidential - Company Proprietary

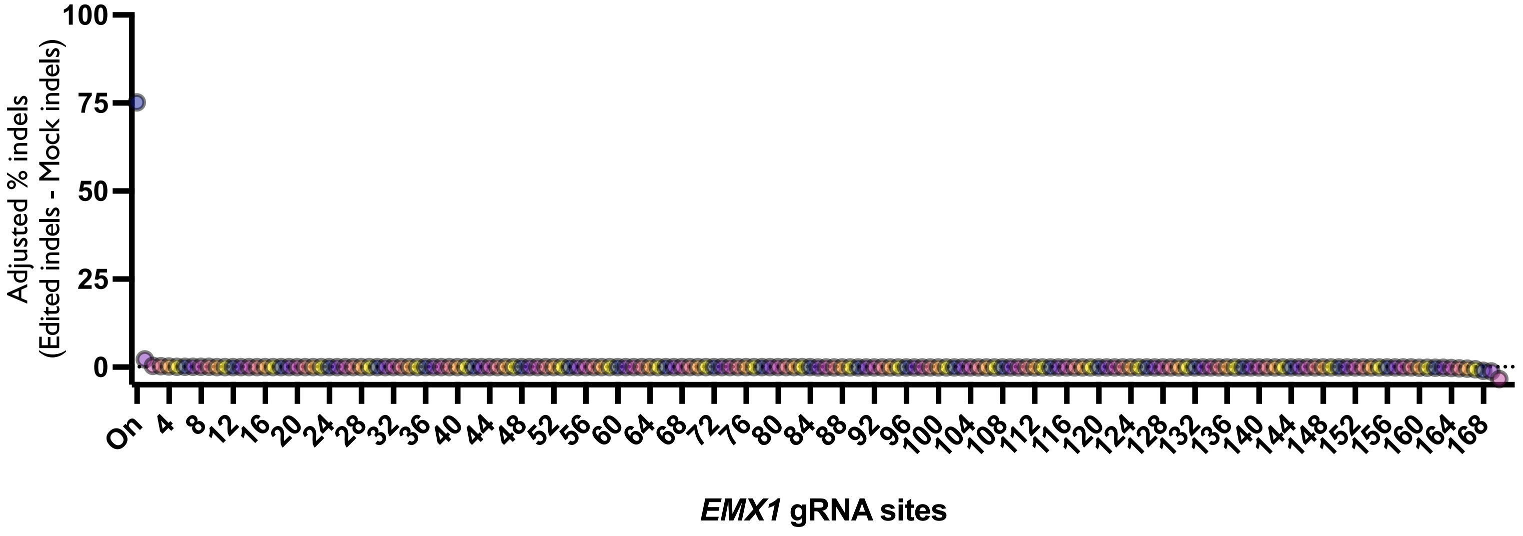


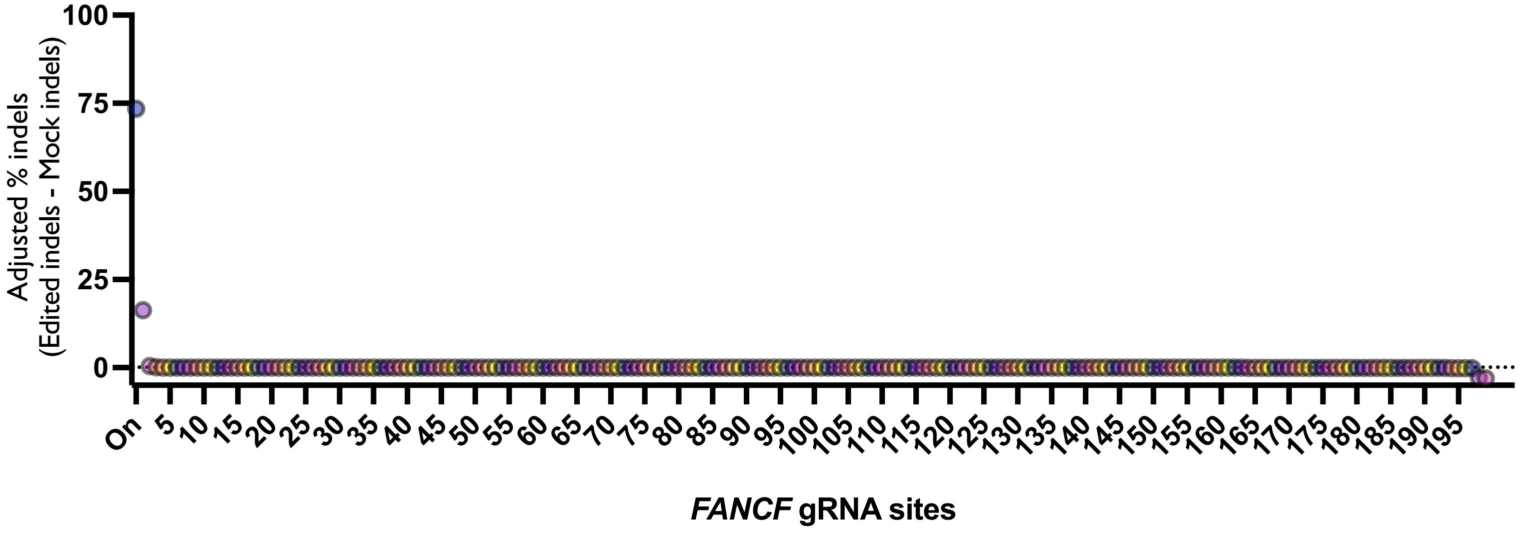


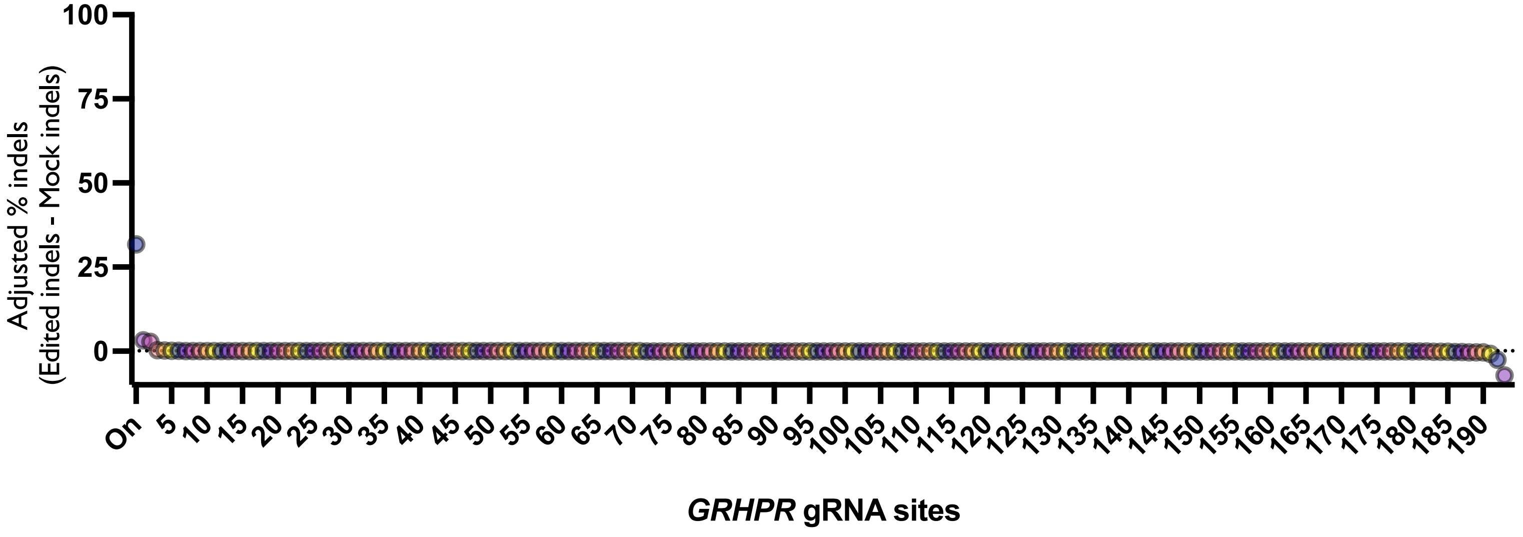


Confidential - Company Proprietary

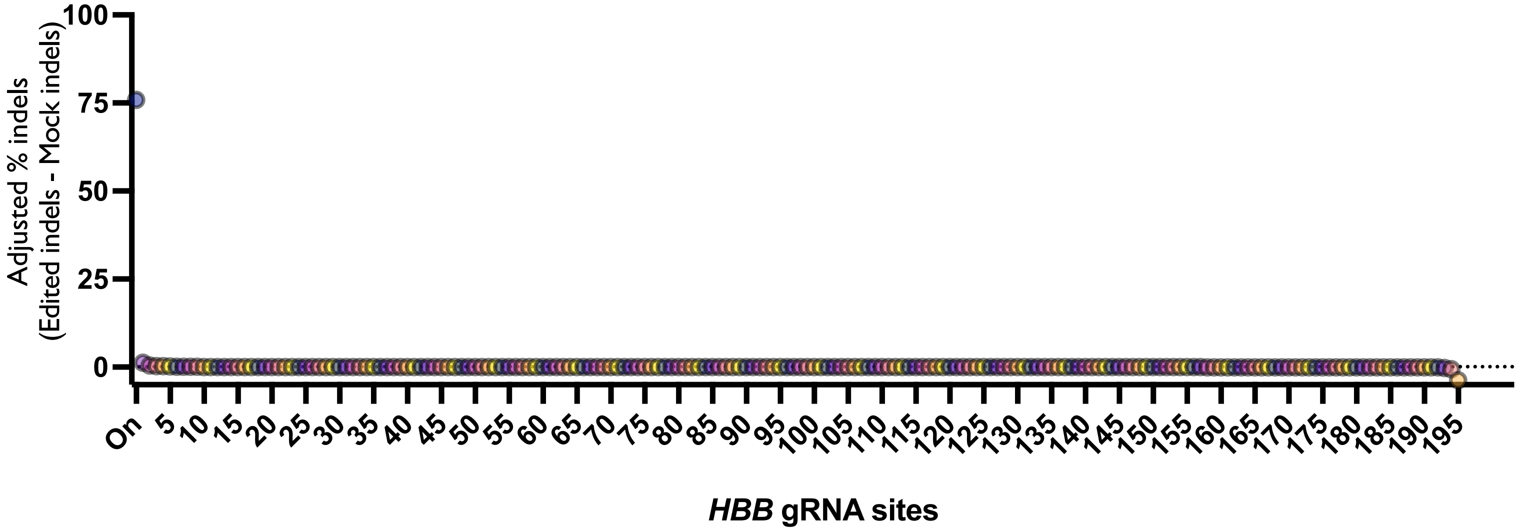


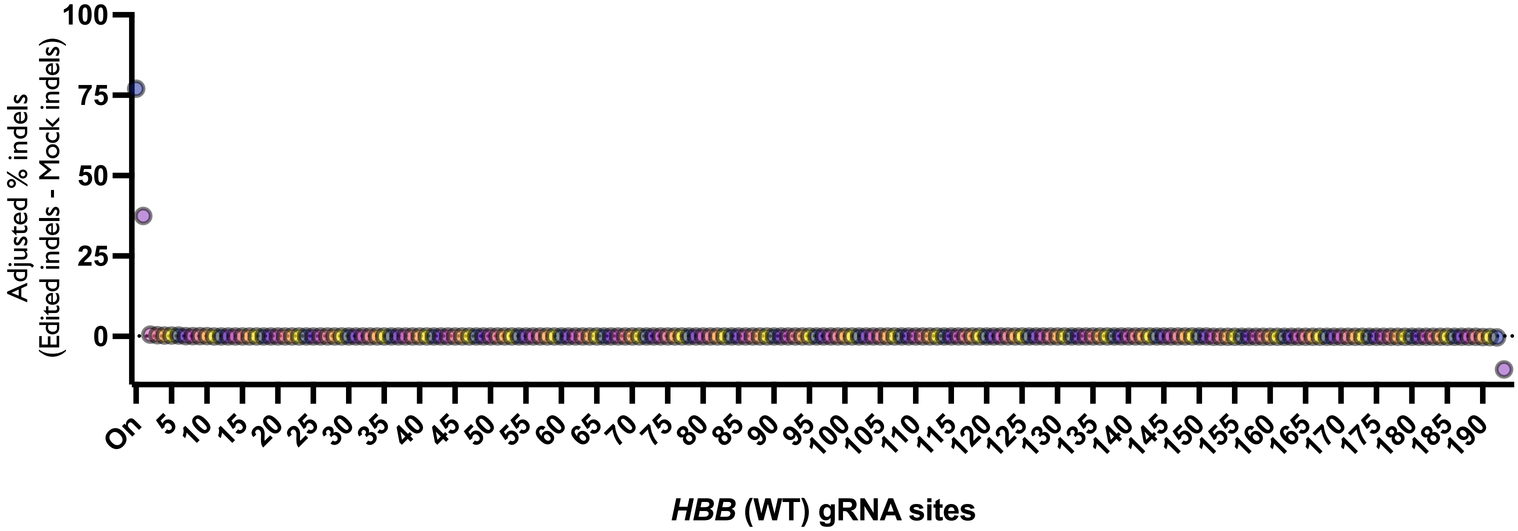


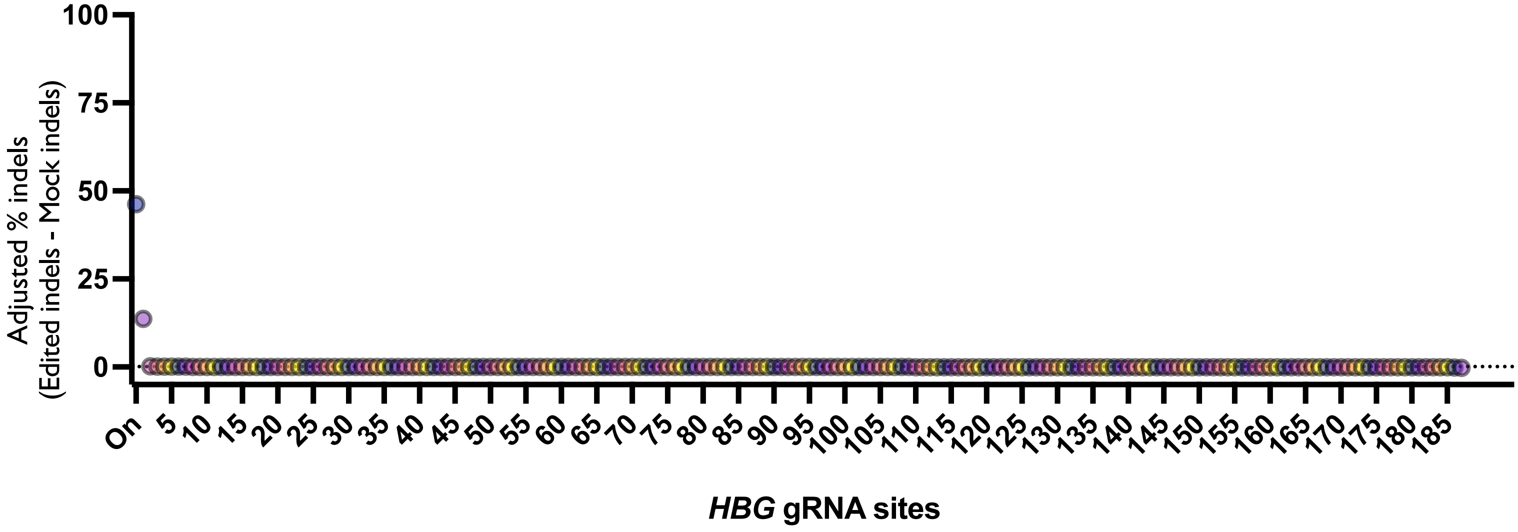


Confidential - Company Proprietary

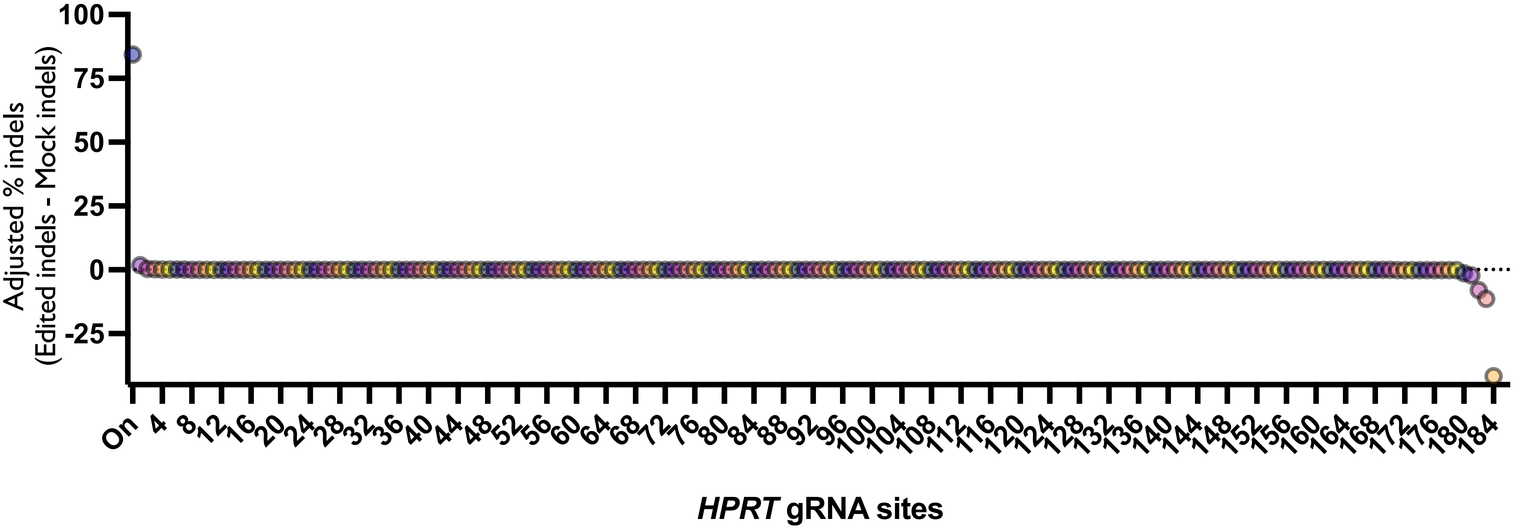


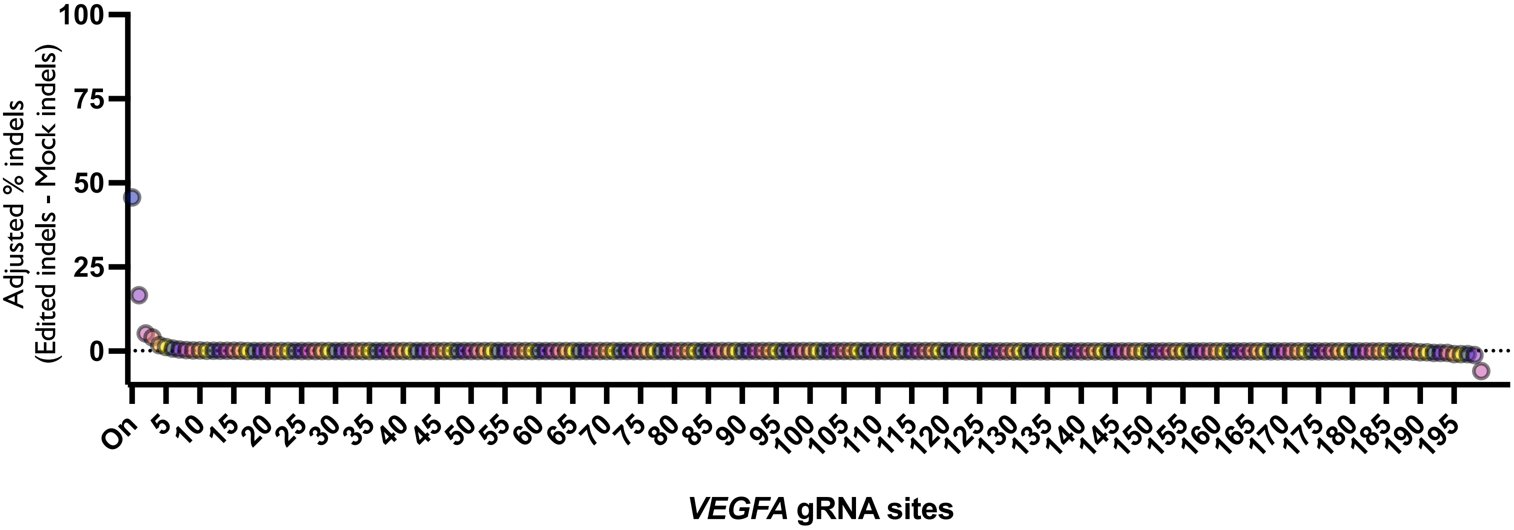


Each graph represents average % indels (Edit-Mock) across donors for each site on panel prior to filtering. Unless otherwise noted, treatments are using HiFi Cas9 and a 20-nt spacer. Dotted line depicts 0.1% indel detection threshold after Mock is subtracted from Edited treatments.

Confidential - Company Proprietary

Supplemental Figure 3: Indel spectrum at on-target site for each gRNA

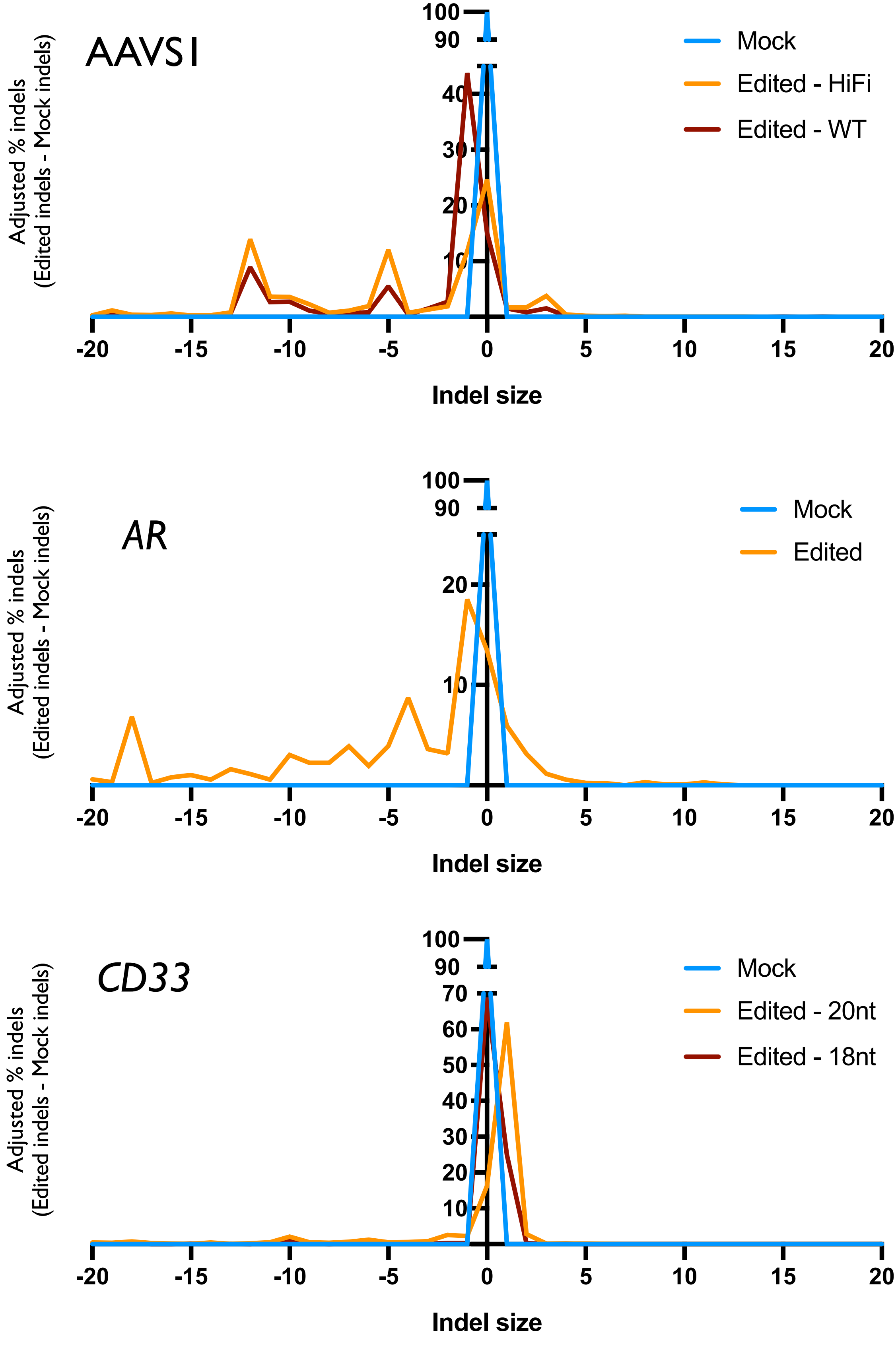


Confidential - Company Proprietary

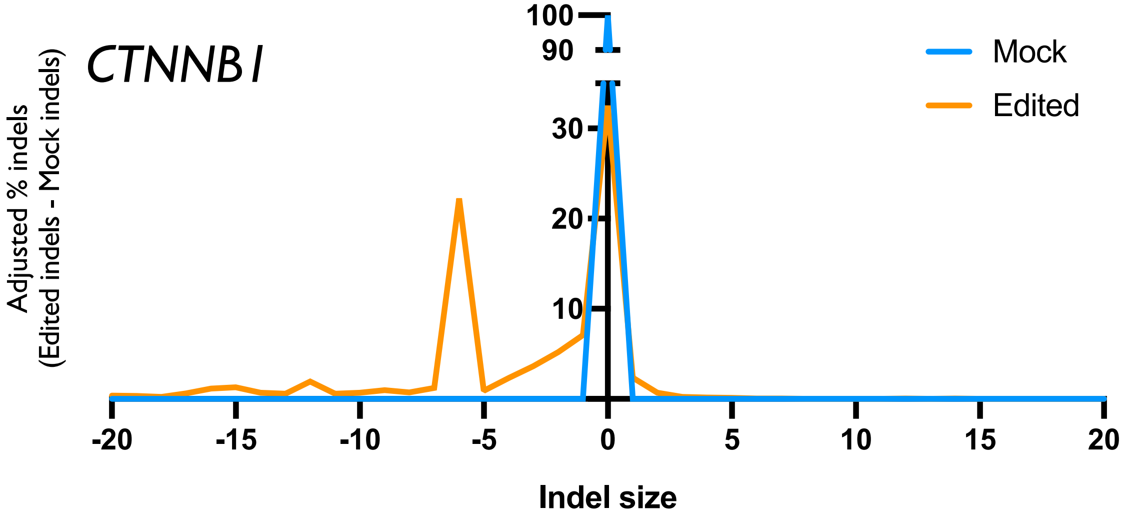


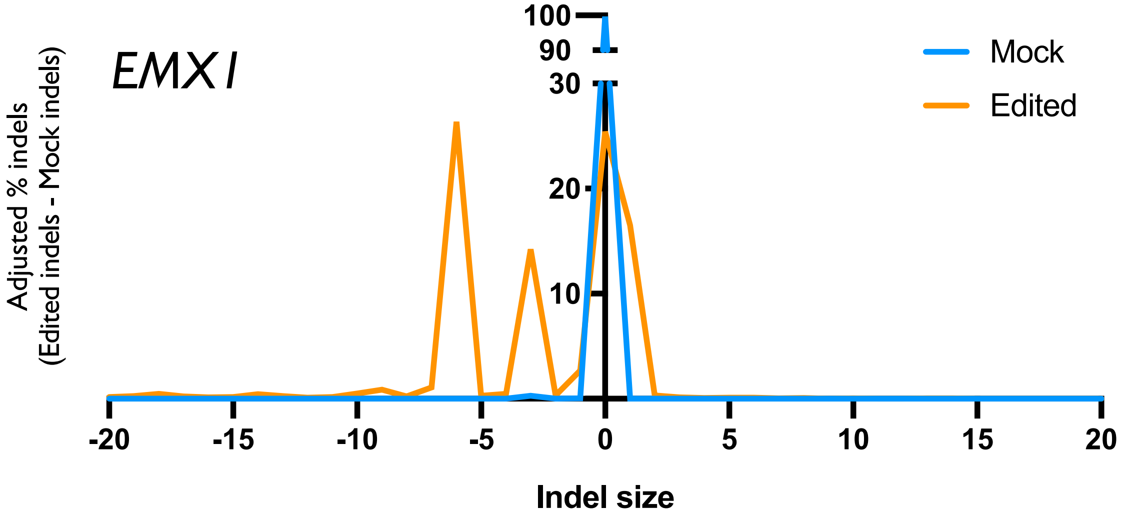


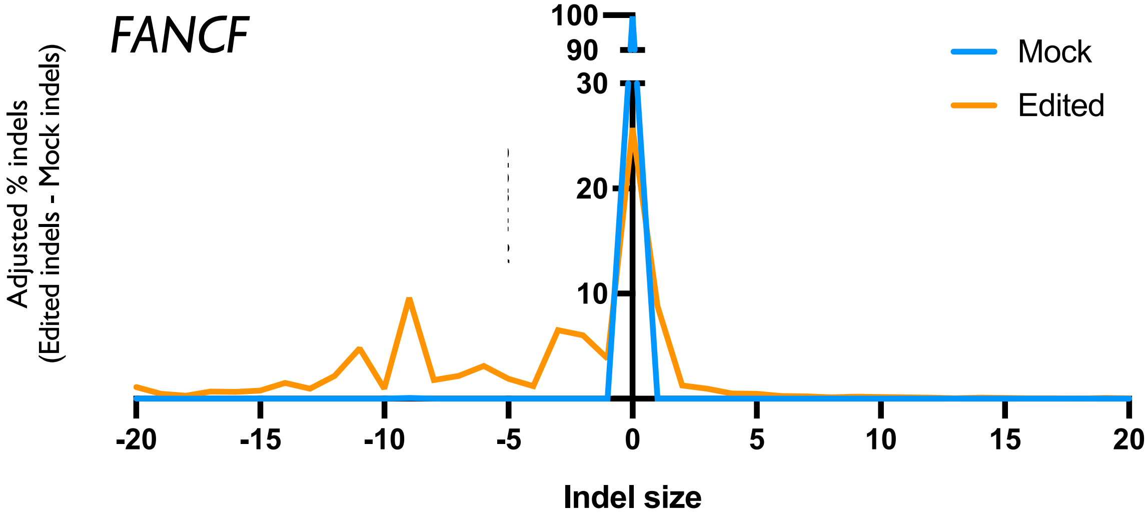


Confidential - Company Proprietary

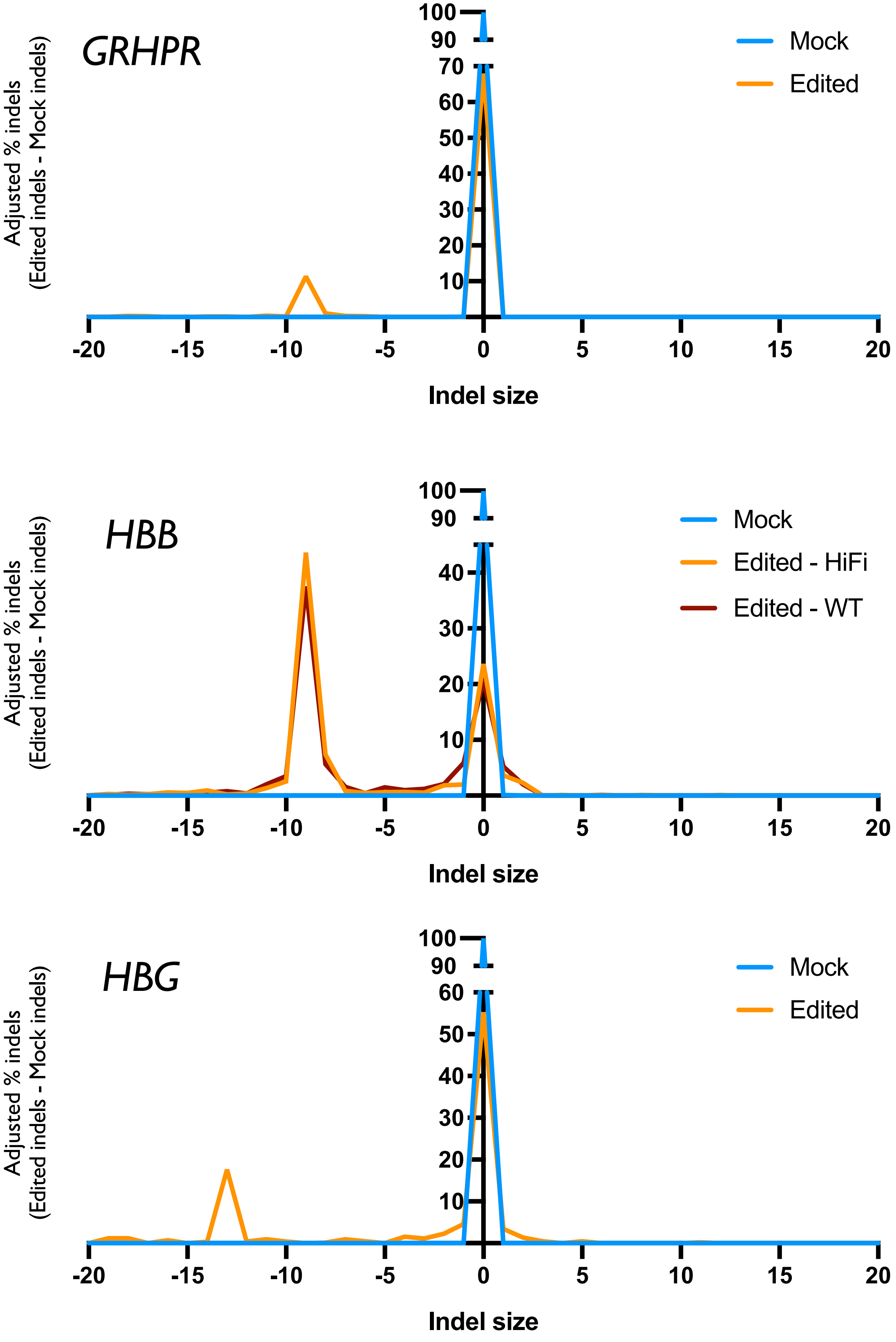


Confidential - Company Proprietary

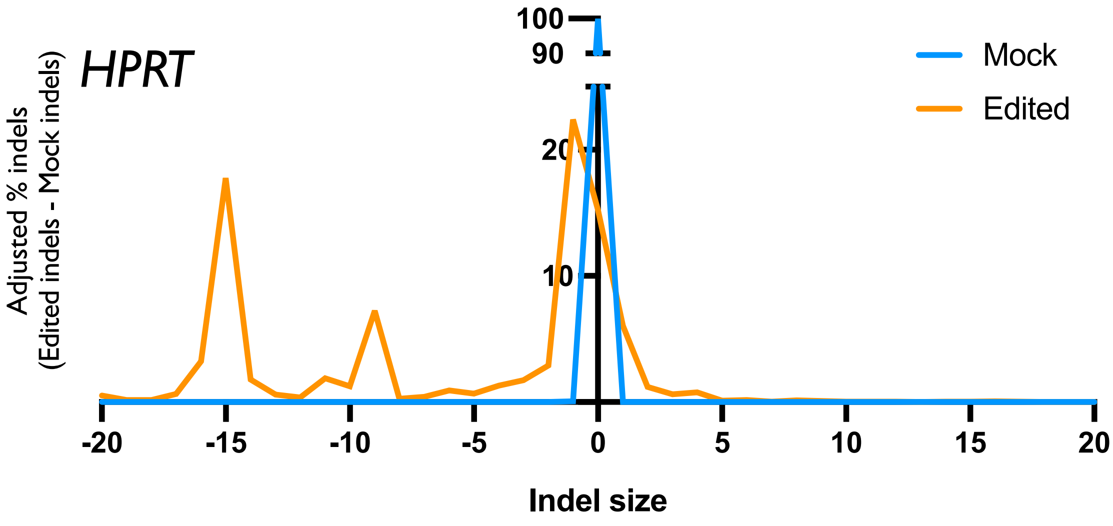


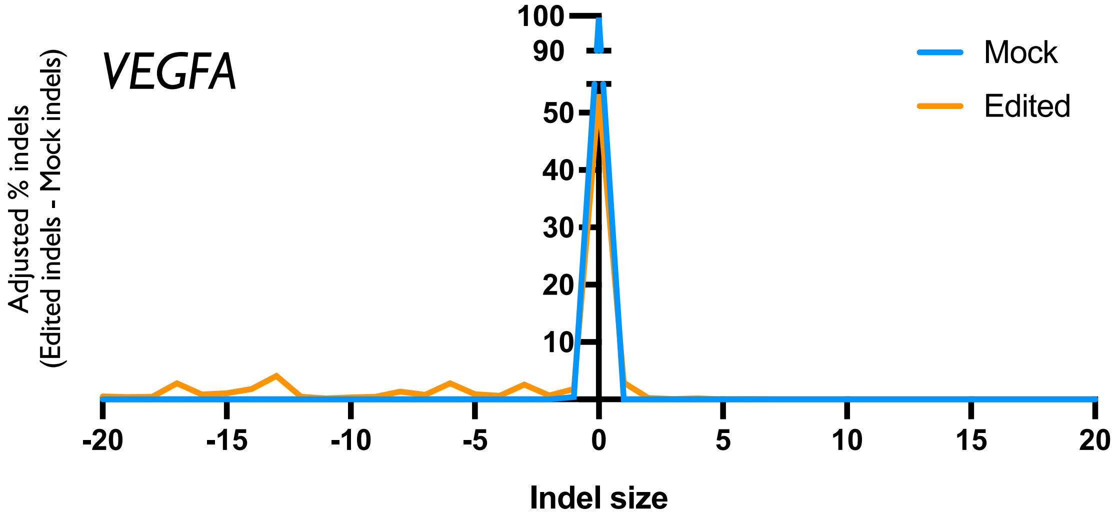


Lines indicate average % indels at each basepair from predicted cut site across 3 donors for Mock, Edited, and Differential (WT Cas9 and 18-nt spacer) treatments. Negative values indicate deletions, positive values indicate insertions.

Supplemental Figure 4: Tool performance summary

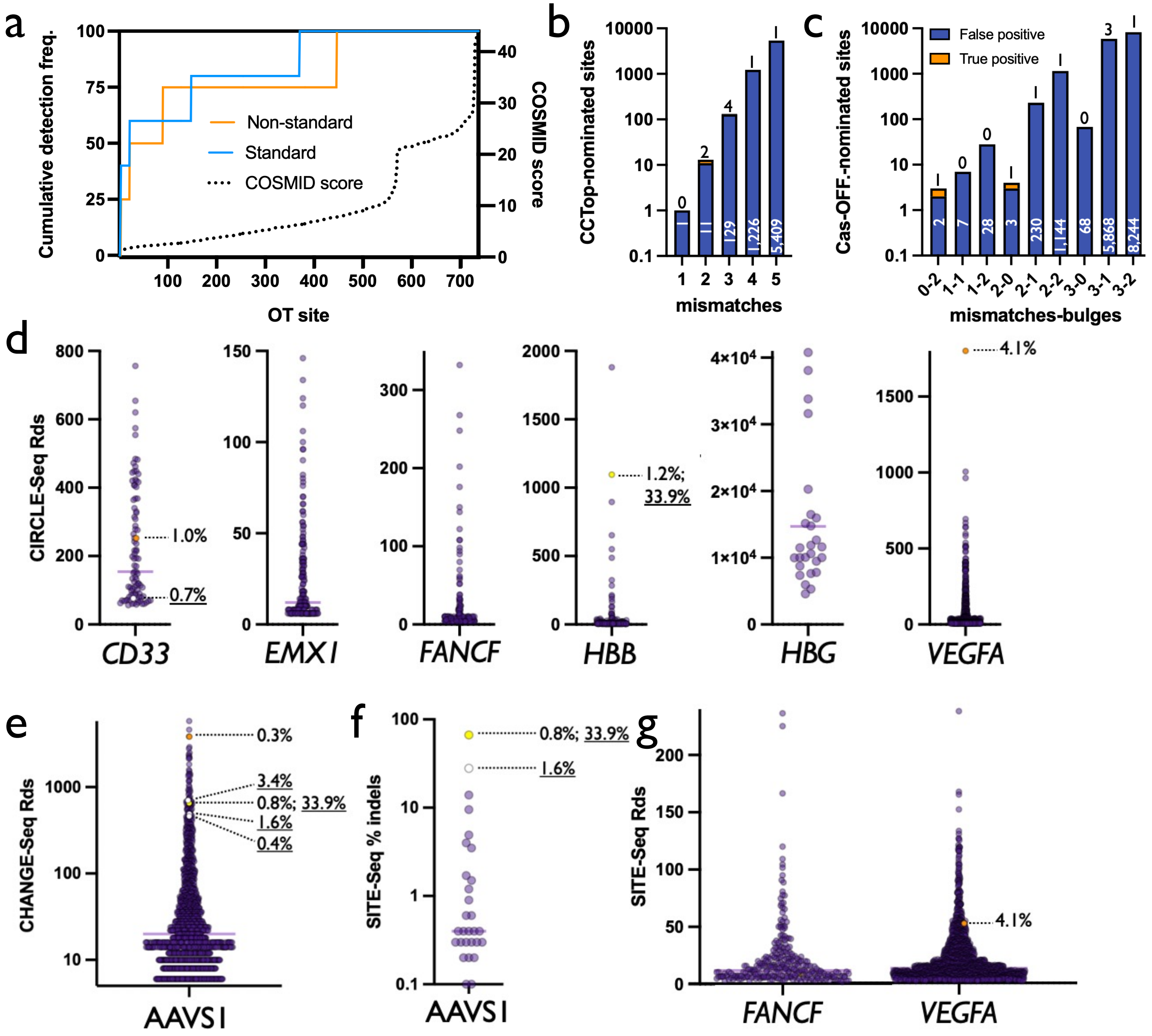


a) For each OT site—rank-ordered left to right from low COSMID score to high along the x-axis—COSMID score and cumulative detection frequency is plotted in standard and non-standard treatments.

b) Each column displays number of true and false positives nominated by CCTop in each bin for a given number of mismatched bases from the target sequence. Shown on base 10 logarithmic scale.

c) Each column displays number of true and false positives nominated by Cas-OFFinder in each bin for a given number of mismatched bases and bulge sites from the target sequence. Shown on base 10 logarithmic scale.

d) Each dot depicts the number of reads covering a single CIRCLE-Seq OT site following editing using the corresponding gRNAs. True positives and corresponding indel frequencies are shown by dotted lines. Orange dots indicate OTs generated by HiFi Cas9, white dot indicates OT generated by WT Cas9, and yellow dot indicates OT generated by both.

e) Each dot depicts the number of reads covering a single CHANGE-Seq OT site following editing using AAVS1 gRNA. True positives and corresponding indel frequencies are shown by dotted lines. Orange, white, and yellow dots indicate OTs generated by HiFi, WT, and both HiFi and WT Cas9, respectively.

f) Each dot depicts % indels at each SITE-Seq OT site following editing using AAVS1 gRNA. True positives and corresponding indel frequencies are shown by dotted lines. Orange, white, and yellow dots indicate OTs generated by HiFi, WT, and both HiFi and WT Cas9, respectively.

g) Each dot depicts the number of reads covering a single SITE-Seq OT site following editing using the corresponding gRNAs. True positive and corresponding indel frequency is shown by dotted line. Orange dots indicate OTs generated by HiFi Cas9.

Confidential - Company Proprietary

Supplemental Figure 5: Likelihood of OTs falling within exons

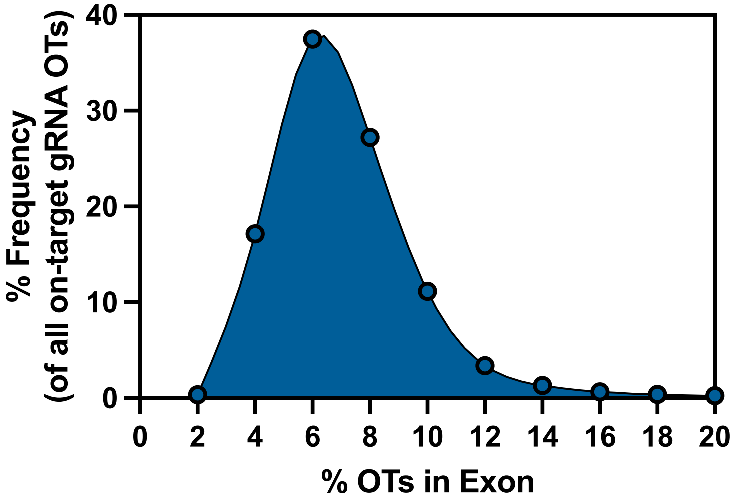


A list of all nominated OT sites for all possible gRNAs targeting 19,222 human genes was generated by the IDT CRISPR-Cas9 Guide RNA Design Checker tool and the frequency of OT sites residing in exons was quantified. Frequencies were binned around 2, 4, 6, 8, 10, 12, 14, 16, 18, and 20 and frequency of OTs residing in exons was plotted accordingly.

Confidential - Company Proprietary
